# Genomic characterization of a nematode tolerance locus in sugar beet

**DOI:** 10.1101/2023.06.22.546034

**Authors:** Katharina Sielemann, Boas Pucker, Elena Orsini, Abdelnaser Elashry, Lukas Schulte, Prisca Viehöver, Andreas E. Müller, Axel Schechert, Bernd Weisshaar, Daniela Holtgräwe

## Abstract

**Background:** Infection by beet cyst nematodes (BCN, *Heterodera schachtii*) causes a serious disease of sugar beet, and climatic change is expected to improve the conditions for BCN infection. Yield and yield stability under adverse conditions are among the main breeding objectives. Breeding of BCN tolerant sugar beet cultivars offering high yield in the presence of the pathogen is therefore of high relevance.

**Results:** To identify causal genes providing tolerance against BCN infection, we combined several experimental and bioinformatic approaches. Relevant genomic regions were detected through mapping-by-sequencing using a segregating F2 population. DNA sequencing of contrasting F2 pools and analyses of allele frequencies for variant positions identified a single genomic region which confers nematode tolerance. The genomic interval was confirmed and narrowed down by genotyping with newly developed molecular markers. To pinpoint the causal genes within the potential nematode tolerance locus, we generated long read-based genome sequence assemblies of the tolerant parental breeding line Strube U2Bv and the susceptible reference line 2320Bv. We analyzed continuous sequences of the potential locus with regard to functional gene annotation and differential gene expression upon BCN infection. A cluster of genes with similarity to the *Arabidopsis thaliana* gene encoding nodule inception protein-like protein 7 (NLP7) was identified. Gene expression analyses confirmed transcriptional activity and revealed clear differences between susceptible and tolerant genotypes.

**Conclusions:** Our findings provide new insights into the genomic basis of plant-nematode interactions that can be used to design and accelerate novel management strategies against BCN.

## 1 Background

Sugar beet is one of the most important crops in the northern hemisphere and contributes about 20% to world-wide sugar production. The ancestor of cultivated sugar beet is the sea beet *B. vulgaris* subsp. *maritima*. White Silesian Beet, a beet segregating in the F2 from a cross of fodder beet and chard, provided the narrow genetic base for today’s sugar beet breeding [1]. An intense focus on yield led to a strong domestication bottleneck [2].

Among the economically most important pests of sugar beet (*Beta vulgaris* subsp. *vulgaris*) is the beet cyst nematode (BCN, *Heterodera schachtii*). Upon infection, the nematodes induce the formation of feeding structures in the roots of the host plant, so called syncytia [3,4]. The development of these syncytia is initiated by secretions from the nematode and expression of specific host plant genes, including expansins, cellulases and endo-1,4-β-glucanases [3–5]. Also, the transcriptome of sugar beet and BCN in compatible and incompatible interactions was studied using RNA-Seq, providing molecular insights into plant-nematode interactions [6]. The harmful effect of the nematode *H. schachtii* is based on nutrient competition and disturbances in the root system of the host plant, which lead to severe growth depression and yield reduction up to 60% [7]. Since growth of *H. schachtii* is supported by increased soil temperatures during the main vegetation period [8], the growth conditions for *H. schachtii* are expected to improve as a result of global warming. This results in an increased yield risk for sugar beet. The use of pathogen resistant or tolerant elite varieties contributes significantly to improved sustainability of sugar production through yield stability. Such varieties address agricultural and social demands on both conventional and organic sugar beet cultivation. Chemical control of BCN by soil decontamination is not possible [9]. Resistant sugar beet varieties that do not allow *H. schachtii* to reproduce during the cultivation phase are available on the market, but they have the disadvantage of penalized yield. In addition, the single gene-based resistances of the sugar beets can be broken relatively quickly by the nematodes and there is a risk of pathotype formation in *H. schachtii* itself [10]. On the contrary, nematode-tolerant beet varieties do not react as strongly with yield depressions when infested with *H. schachtii* [11] and therefore represent an economically remunerable trait to breed for. From a genetic point of view, nematode tolerance is a quantitative resistance that inhibits BCN development [11,12] in terms of both quantity and quality.

Molecular genetic markers associated with BCN resistance or tolerance are an important step towards the identification of trait-associated genes [13]. The first known nematode resistance gene, *Hs1^pro-1^*, encodes a protein harboring a leucine-rich domain. *Hs1^pro-1^*was introduced into sugar beet as part of a translocation from the crop wild relative *Patellifolia procumbens* [9]. Since then, other trait-associated genomic regions have been identified, including a region conferring nematode tolerance from sea beet [14]. Stevanato *et al.* [13] identified a region on chromosome 5 (chr5) of the sea beet genotype WB242 (*BvmHs^-1^*) and published a molecular marker, designated SNP192, linked to nematode tolerance. Using segregation analyses, the group was able to show the monogenic inheritance of the trait. However, the region was not further defined or described, and variation for the trait nematode tolerance was observed by breeders although SNP192 was homozygous in the genotypes studied.

To gain gene-level information for traits of interest, genome sequences of accessions harboring these traits and comprehensive annotations are needed. In combination with methods like mapping-by-sequencing (MBS), this allows a detailed investigation of agronomically important regions. With regard to sugar beet, high-quality genome sequences are available for two different accessions. These are the inbred line EL10 [15] and the ‘reference genotype’ KWS2320 (referred to here as ‘2320Bv’) with the sequence identifier Refbeet-1.0 [16]. An improved version that, among other data, also incorporates the genetic map BeetMap-3 [17] is publicly available with the identifier RefBeet-1.5 (https://jbrowse.cebitec.uni-bielefeld.de/RefBeet1.5/). In addition, draft genome assemblies of *B. patula* and *B. vulgaris* subsp. *maritima* WB42 [18] that represent crop wild relatives of sugar beet, are publicly available. No high-quality genome sequence of a BCN tolerant breeding line was available until now, and short read assemblies are not suitable for MBS approaches and subsequent detailed trait locus analysis.

Access to BCN tolerant sugar beet cultivars that produce high yield even on BCN containing soil is of high relevance for sugar production in the northern hemisphere. In this study, we targeted the nematode tolerance (NT) locus on chr5 with genomic approaches to further delimit this locus and to describe the genes included. High-continuity genomic resources of the tolerant genotype Strube U2Bv and the susceptible sugar beet reference genotype 2320Bv were developed. MBS of tolerant and susceptible lines to BCN was used together with RNA-Seq data generated from infection experiments. This allowed to further characterize the potential genomic locus responsible for nematode tolerance (NT) in sugar beet. Our results will benefit breeding approaches and enable a better control of the yield-diminishing BCN disease.

## 2 Results

### 2.1 Generation and phenotyping of a segregating F2 population

The mapping population STR-NT was derived from the nematode susceptible maternal line Strube U1Bv and the tolerant paternal line Strube U2Bv. A total of 406 F2:3 lines segregated for tolerance to BCN with continuous variation. The distribution of adjusted means of quantified tolerance per single plant ranged from -0.105 to 16.835, corresponding to 0 to 280 cysts counted per plant (Figure 1A). Twenty tolerant F2:3 lines with low numbers of cysts and 16 susceptible F2:3 lines with high number of cysts were identified as extremes of the phenotypic distribution. Analysis of coefficient of variation (CV) suggested no further segregation for NT on these families so that their further investigation was performed through MBS.

**Figure 1:**
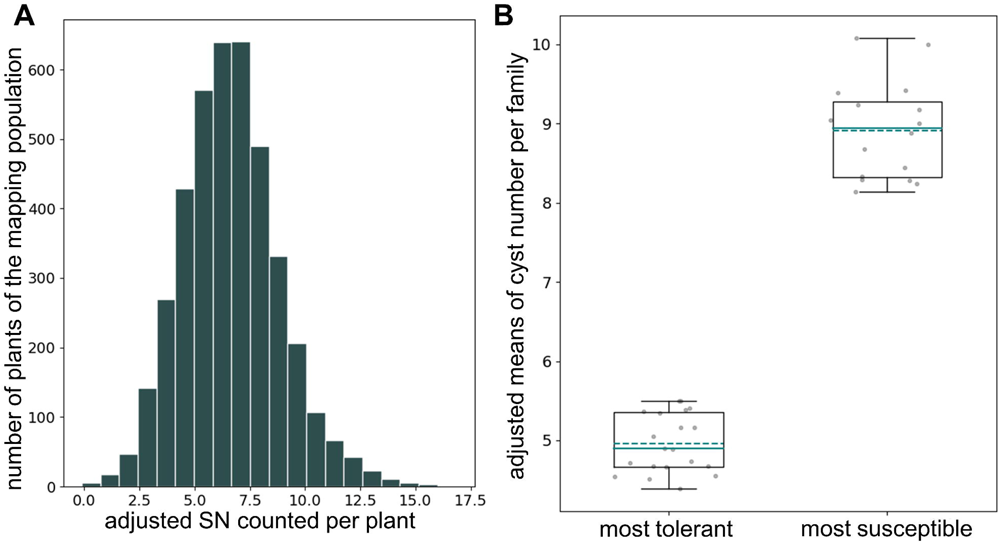
Phenotyping of the population STR-NT segregating for BCN tolerance. **A)** Histogram of adjusted cyst counts (SN) of single plants. SN = squared root of the number of cysts counted per plant (defined as in the methods section). **B)** Boxplot of the most tolerant and most susceptible F2-lines represented as families chosen for MBS. Adjusted means of the most tolerant families range from 4.39 to 5.5 corresponding to 19-30 cysts. Adjusted means of the most susceptible families range from 8.14 to 10.08 corresponding to 66-101 cysts.

Adjusted means per line (Figure 1B, Additional file S2A) were used for QTL mapping together with the genotypic data of 194 KASPar markers. A major QTL with an additive effect corresponding to 11 cysts was detected on the north of chr5 at 10 cM. This QTL explained 23% of the phenotypic variance. The limit of detection (LOD) support interval spanned the region from 9 to 13 cM between the new marker BR1180 and SNP192. The tolerant allele was derived from the nematode tolerant parent Strube U2Bv. No dominance effects were detected, heterozygous lines display an intermediate phenotype between tolerant and susceptible lines.

For the MBS approach, the nine most tolerant lines were represented in individual libraries and the eleven ‘second best’ tolerant lines were combined and sequenced in the “pool low”. The opposite phenotypic extreme included 16 susceptible lines. The nine most susceptible lines were used to create individual libraries, whereas the remaining seven lines were combined in the “pool high” and sequenced. For the tolerant lines, a total of 244 Gbp of Illumina reads totaling a calculated 20.4x genome coverage were obtained, whereas for all susceptible lines, 189 Gbp of reads with an estimated genome coverage of 19.8x were reached.

### 2.2 Assembly and annotation of genome sequences of the nematode-tolerant parent Strube U2Bv and the reference genotype 2320Bv

High assembly continuity for both, the nematode tolerant parent Strube U2Bv and the susceptible reference genotype 2320Bv, is indicated by the N50 values (Table 1). The assembly sizes of 596 Mbp (U2BvONT) and 573 Mbp (2320BvONT) exceed the size of RefBeet-1.5 by 29 Mbp and 6 Mbp, respectively. RepeatMasker [19] results indicated that both assemblies have a repeat content of more than 65%.

**Table 1:** Assembly statistics of U2BvONT and 2320BvONT v1.0. Assembly size, number of contigs, N50 values, BUSCO completeness, repeat content, number of predicted genes, and number of predicted mRNAs are shown.

|  | 2320BvONT v1.0 | U2BvONT v1.0 |
| --- | --- | --- |
| Assembly size [bp] | 573,025,584 | 596,437,702 |
| Sequences | 79<br>(9 pseudochromosomes + 70<br>contigs larger than 100 kb) | 155<br>(9 pseudochromosomes + 146<br>contigs larger than 100 kb) |
| N50 [bp] | 54,419,778 | 54,373,962 |
| BUSCOs (n=2326) | C:93.7% [S:92.1%, D:1.6%],<br>F:3.0%, M:3.3%<sup>*</sup> | C:93.6% [S:92.0%, D:1.6%],<br>F:3.2%, M:3.2%<sup>*</sup> |
| Repeat content | 65.21% | 65.85% |
| Number of<br>predicted genes | 27,840 | 28,871 |
| Number of<br>predicted mRNAs | 36,350 | 36,728 |

<sup>\*</sup>Abbreviations: C = Complete, S = Single-copy, D = Duplicated, F = Fragmented, M = Missing

The initial assemblies of Strube U2Bv and 2320Bv prior to the scaffolding process comprised 206 and 129 sequences, respectively. In total, 87.0% total bases of U2BvONT, represented by 60 initial contigs, were anchored to nine pseudochromosomes using Beet-Map3 markers, whereas only a slightly higher percentage of bases (88.3%) were genetically anchored in 2320BvONT represented by 59 initial contigs. In summary, 1,237 markers anchored the 2320BvONT assembly, whereas 1,238 markers anchored the scaffolds in the U2BvONT assembly. A new set of 187 markers was designed based on data for Strube U2Bv and successfully applied to genotyping in the StrUBv F2-population. A subset of 165 markers confirmed the co-linearity of both assemblies and with Beet-Map3 (Additional file S1A). However, these markers did not always allow exact genetic anchoring because of missing information regarding the orientation of contigs, therefore such ambiguous contigs were not placed in the pseudochromosomes.

Synteny analysis between 2320BvONT and RefBeet annotations revealed a mRNA-based synteny of 97.3% (35,363) with a depth of 1 indicating a high similarity. Duplicated mRNAs in 2320BvONT have, in comparison to RefBeet, a proportion of 0.6% (231), and 2.1% (756) of 2320BvONT mRNAs were not found to be syntenic with any RefBeet mRNAs.

Investigation of the synteny between 2320BvONT and U2BvONT showed a 1:1 relation of 98.3% (35,721) of all mRNAs. Duplicated mRNAs in 2320BvONT occur, in comparison to U2BvONT, with 0.6% (200); 1.2% (429) have no syntenic anchor in U2BvONT. Genes without a syntenic counterpart were investigated to identify potential functions not present in the respective other genome. However, BLAST and InterProScan revealed no unique hit in either of the genomes.

Several large structural variations differentiate the genomes of 2320BvONT and U2BvONT. In total, 63 Mbp, divided into 88 different regions, are inverted between both genomes. The largest inversion on chr3 with a size of approximately 21 Mbp spans almost three contigs. Additionally, more than 3,000 translocations with a combined size of approximately 33 Mbp were identified (Additional file S2B).

### 2.3 NT locus detection through mapping-by-sequencing

A total of 2,049,007 variant positions were identified by comparing variants identified as homozygous in both parents and as heterozygous in the F1. The average delta allele frequency (dAF) in 10 SNP windows across all chromosomes is approximately 0.103 (±0.079). A dAF > 0.5 in a 10 SNP window was only detected on chr5 (Figure 2A) and on no other chromosome or contig of the U2BvONT assembly. The dAF plots of all nine chromosomes are provided in Additional file S2C. The borders of our potential locus of interest were restricted by the occurrence of a dAF > 0.5 throughout five consecutive 10 SNP windows. This 50 SNP window delimitation resulted in a genomic interval ranging from position 452,859 bp – 4,557,625 bp on chr5 of the U2BvONT assembly.

**Figure 2:**
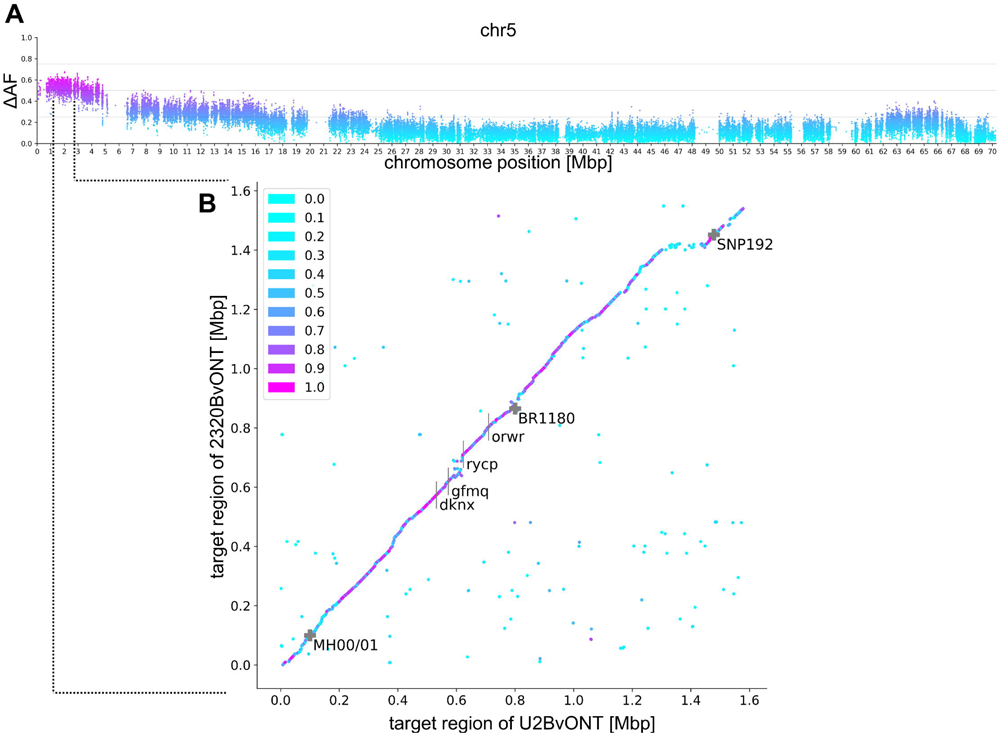
Boundaries of the potential NT locus region. **A)** Delta allele frequency (ΔAF) distribution on chr5 of the U2BvONT assembly. Higher dAF, purple color; lower dAF, light blue. **B)** Dot plot comparing the potential NT locus region in U2BvONT and 2320BvONT. Magenta and purple dots represent high BLAST identity between both genome sequences. The potential NT locus was further delimited by the genetic markers MH00/01 and BR1180. Marker SNP192 [13] is linked but located further to the south of the chromosome. The 4-letter codes represent gene IDs (see text).

This clear interval of about 4 Mbp from MBS was further restricted using marker analyses on a few extreme F2 genotypes and the established susceptible and tolerant standard lines. The genetic markers MH00/01, BR1180 and additional flanking markers including SNP192 enabled, by graphical genotyping of recombination events in the phenotyped lines, a further containment of the potential NT locus (Additional file S2D). The size was restricted to about 0.7 Mbp with coordinates 1,321,396 – 2,021,946 bp in U2BvONT and 1,389,131 – 2,154,734 bp in 2320BvONT. The published marker SNP192 is located further south on chr5 at position 2,700,363 bp in U2BvONT and 2,741,678 bp in 2320BvONT (Figure 2B). The sequences of the whole region are continuous in both assemblies.

### 2.4 Characterisation of the potential NT locus

Overall, very high synteny was detected within the marker-restricted potential NT locus in U2BvONT and 2320BvONT (Figure 2). However, reduced synteny was observed between the genes gfmq and rycp. Within this region, we identified a cluster of genes (Figure 3) with similarity to *AtNLP7*. Analysis of the direct target genes of the AtNLP7 TF revealed that genes annotated with ‘response to nematode’ are significantly overrepresented among the targets (Additional file S1B). Several other terms possibly related to BCN infection, like ‘response to stress’, ‘signal transduction’, and ‘response to other organism’ were found to be overrepresented as well. Therefore, we focused on this region, which is smaller than the marker-restricted potential NT locus and call it ‘functionally restricted potential NT locus’.

**Figure 3:**
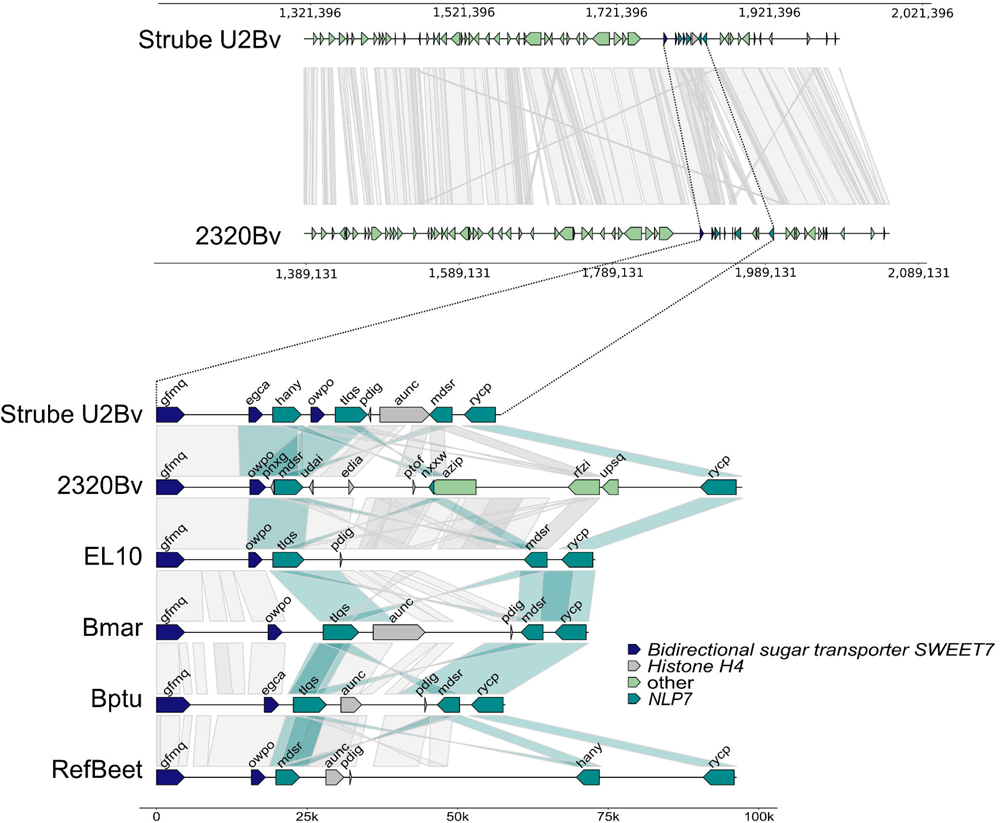
Overview of the potential NT locus (top) and illustration of the functional restriction of the potential NT locus (bottom). Each line represents the sequence region between the genes gfmq and rycp for the respective genotype/species (*B. vulgaris* subsp. *vulgaris*: U2BvONT (Strube U2Bv); 2320BvONT (2320Bv); EL10; *B. vulgaris* subsp. *maritima* WB42 (Bmar), *B. patula* BETA548 (Bptu), RefBeet 1.5 (RefBeet)). The grey areas connecting the sequences indicate synteny. Some connecting areas are highlighted in turquoise to show synteny between the genes with similarity to *AtNLP7*. *Bidirectional sugar transporter SWEET7*: gfmq, egca, owpo; *Histone H4*: pdig, aunc, udai, edia, ptof; *BvNLP7*: hany, tlqs, mdsr, rycp, nxxw; other: azip, rfzi, upsq.

The number of *BvNLP7* genes differs in both assemblies. In total, four and three genes are functionally annotated as ‘NLP7’ in the U2BvONT and 2320BvONT genome assembly, respectively. All of them are located within the delimited potential NT locus (Figure 3, Table 2). A manual check of reading frames in the region of both assemblies revealed no additional *NLP7*-like sequence.

**Table 2:** Gene/allele names for the *NLP7*-like genes. The short name is used for more clarity in the text. T = tolerant genotype (Strube U2Bv); S = susceptible genotype (2320Bv).

| Short name | Genotype | Gene Identifier |
| --- | --- | --- |
| *NLP7-T1* | Strube U2Bv | Bv05_g11459_hany.t1 |
| *NLP7-T2* | Strube U2Bv | Bv05_g11464_mdsr.t1 |
| *NLP7-T3* | Strube U2Bv | Bv05_g11461_tlqs.t1 |
| *NLP7-T4* | Strube U2Bv | Bv05_g11465_rycp.t1 |
| *NLP7-S1* | 2320Bv | Bv05_g11045_mdsr.t1 |
| *NLP7-S2* | 2320Bv | Bv05_g11050_nxxw.t1 |
| *NLP7-S3* | 2320Bv | Bv05_g11053_rycp.t1 |

The genes with similarity to *AtNLP7* were compared in multiple sequence alignments at coding sequence (CDS) and amino acid (aa) sequence level (Additional file S3, Additional file S4, Additional file S5, Additional file S6, Additional file S1C).

The CDS of *NLP7-T4* and *NLP7-S3* show an identity of 99.5% (Additional file S1C) with only 15 single nucleotide variant positions. At the aa level, these encoded proteins differ only at two amino acid positions, namely isoleucine to valine and asparagine to serine (I878V and N927S in U2BvONT > 2320BvONT). Both conservative exchanges are a result of a single nucleotide variant. Similarly, *NLP7-T2* shows CDS identity to *NLP7-S1* of 93.49% (Additional file S1C). Due to the high identity, we formally consider these gene structures as allelic. *NLP7-S2* is truncated in comparison to *AtNLP7* and codes only for a part of the protein.

All candidate genes were compared to *AtNLP7* via percent identity matrices (Additional file S1C). The comparison was performed for CDS and aa sequences as well as for the PB1 domain. The U2BvONT *NLP7*-like candidate genes show a relatively low sequence identity to *AtNLP7* (Additional file S1C, CDS), ranging from 43.2% (*NLP7-T2*), 44.6% (*NLP7-T1*), and 45.1% (*NLP7-T3*) to 63.5% (*NLP7-T4*). For the three 2320BvONT *BvNLP7* genes, the sequence identities are comparable (44% (*NLP7-S1*), 52.6% (*NLP7-S2*) and 63.5% (*NLP7-S3*)). Next, the key polymorphisms at the described conserved positions (see Background) were investigated (Additional file S2E). The core aa positions of the PB1 domain are completely missing from the sequence of NLP7-T3. K867, D909 and E913 are conserved in all other candidates and D911 is conserved in the candidates except NLP7-S1 and NLP7-T3. The completely conserved residue S205 as well as the PB1 domain are absent from NLP7-S2 due to its truncation. The protein sequences of NLP7-T4 and NLP7-S3 carry S205, and in NLP7-T2, NLP7-T3 and NLP7-S1 this position is substituted by threonine (S205T), another polar amino acid and possible phosphorylation site. Only the protein sequence of NLP7-T1 holds a lysine (S205K).

### 2.5 Tolerant and susceptible genotypes show diverse expression of *BvNLP7* genes

Gene expression was investigated in an infection assay with *H. schachtii*. A principal component analysis (PCA) was conducted to assess the sample distribution (Additional file S2F). Next, differentially expressed genes (DEGs) were identified. A total number of 2,263 genes with a padj < 0.05 was detected to be differentially expressed between all samples of the tolerant and all samples of the susceptible lines (Additional file S1D, Additional file S1E, Additional file S1F). Normalized counts were generated for all U2BvONT genes in tolerant (BR12 and Strube U2Bv) and BCN susceptible lines (Strube U1Bv and SUS3). All four *BvNLP7* genes are expressed in all genotypes under both conditions. Clear differences are visible for the four *BvNLP7* genes. In particular, the two genes *NLP7-T1* and *NLP7-T2* are significantly lower expressed in both susceptible genotypes compared to both tolerant genotypes.

The genes *NLP7-T1* and *NLP7-T2* are upregulated in tolerant genotypes in comparison to susceptible genotypes, independent of the inoculation. In addition, all four *BvNLP7* genes are significantly higher expressed in inoculated samples of the most tolerant F2 line BR12 than in inoculated SUS3 samples (Table 3; Figure 4). Further, both NLP-T1 and NLP-T2 are significantly higher expressed in samples of the inoculated tolerant line U2Bv than in samples of the inoculated susceptible line U1Bv.

**Figure 4:**
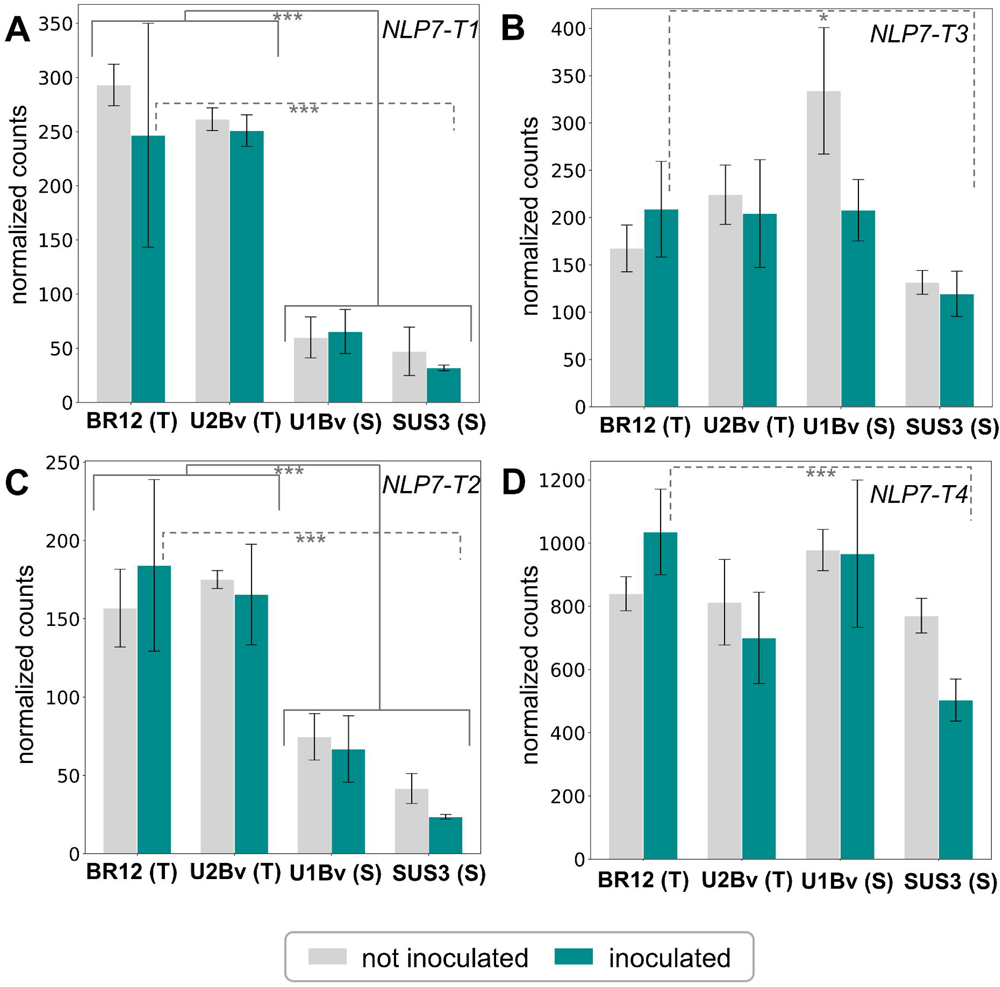
Mean normalized counts (n=3) for the *BvNLP7* genes. RNA-Seq mappings were performed against the U2BvONT assembly. Turquoise bars represent inoculated samples, whereas grey bars show the counts for non-inoculated samples. BR12 and Strube U2Bv are tolerant genotypes, whereas Strube U1Bv and SUS3 are susceptible genotypes. Samples were taken at 21 days post infection (dpi). A) *NLP7-T1*, B) *NLP7-T3*, C) *NLP7-T2*, D) *NLP7-T4*. * = padj < 0.05, ** = padj < 0.01, *** = padj < 0.001.

**Table 3:** Log_2_ fold change (log2FC) and adjusted p-value (padj) of the expression analysis for all *BvNLP7* genes. A positive value for the log2FC indicates higher expression in tolerant genotypes. The comparison ‘all tolerant vs. all susceptible’ involves all samples, inoculated and not inoculated, stratified by the tolerant vs. susceptible genotypes.

| Short name | U2BvONT gene identifier | log2FC | Padj | Comparison |
| --- | --- | --- | --- | --- |
| *NLP7-T1* | Bv05_g11459_hany.t1 | 2.97 | 2.5e-10 | inoculated tolerant line BR12 vs.<br>inoculated susceptible line<br>SUS3 |
| *NLP7-T2* | Bv05_g11464_mdsr.t1 | 2.99 | 2.4e-12 | inoculated tolerant line BR12 vs.<br>inoculated susceptible line<br>SUS3 |
| *NLP7-T3* | Bv05_g11461_tlqs.t1 | 0.82 | 0.027 | inoculated tolerant line BR12 vs.<br>inoculated susceptible line |
|  |  |  |  | SUS3 |
| *NLP7-T4* | Bv05_g11465_rycp.t1 | 1.03 | 6.4e-05 | inoculated tolerant line BR12 vs.<br>inoculated susceptible line<br>SUS3 |
| *NLP7-T1* | Bv05_g11459_hany.t1 | 1.93 | 7.8e-06 | inoculated tolerant line Strube<br>U2Bv vs. inoculated susceptible<br>line U1Bv |
| *NLP7-T2* | Bv05_g11464_mdsr.t1 | 1.29 | 5.3e-04 | inoculated tolerant line Strube<br>U2Bv vs. inoculated susceptible<br>line U1Bv |
| *NLP7-T1* | Bv05_g11459_hany.t1 | 2.34 | 6.5e-27 | all tolerant vs. all susceptible |
| *NLP7-T2* | Bv05_g11464_mdsr.t1 | 1.71 | 2.2e-13 | all tolerant vs. all susceptible |

## 3 Discussion

Phenotypic evaluation of the 406 F2:3 families segregating for BCN tolerance, MBS and generation of highly continuous, well annotated genome sequences allowed characterization of the potential NT locus as well as development of tightly linked molecular markers. This trait region is located further north on chr5 when compared to the published BCN tolerance marker SNP192 [13].

The identification of extremes in the BNC tolerance distribution and the continuous variation of cyst numbers per F2 line demonstrated that the STR-NT population as well as the method for BNC tolerance scoring was suitable for the MBS analysis. Variant detection and dAF calculation were aided by the two new long read-based sugar beet genome sequence assemblies of the susceptible genotype 2320Bv and the BNC tolerant line Strube U2Bv.

In case of monogenic traits, the difference in allele frequency at the causal gene locus should ideally be 100% and thus have a dAF value of 1. This can be achieved for phenotypic data of traits scored without confounding environmental and/or technical effects. An example is the the *RED* locus for sugar beet hypocotyl color which is classified as either “green” or “red” [20]. The potential NT locus shows up in MBS as a single locus in the north of chr5, but the locus reflects a major QTL that independently does not fully explain tolerance to BCN in Strube U2Bv. Therefore, dAF values at the NT locus are expected to be lower than 1.

Although a candidate interval on chr5 was already visible in a preliminary analysis when using the initial RefBeet sequence [16], the data quality was substantially improved in the current study. In fact, the new assemblies 2320BvONT and U2BvONT resolve previously collapsed or duplicated regions and offer continuous sequences for the region of interest on chr5.

The *B. vulgaris* repeat content was previously estimated to be 63% [21]. The new assembly 2320BvONT contains a comparable proportion of repeats, whereas RefBeet contains 42.3% of repeats [16]. This indicates that 2320BvONT contains a substantial amount of sequences not present in the RefBeet assembly. The U2BvONT assembly, which displays similarly high quality and represents the first full assembly of a BCN tolerant line, will serve as an additional resource for genome-informed sugar beet breeding.

The slightly higher number of genes predicted for the U2BvONT assembly can be explained by the ∼20 Mbp larger assembly size. This additional sequence possibly originates from remaining heterozygosity in Strube U2Bv or may be partly explained by a slightly higher repeat content. The *ab initio* gene prediction for both genome sequences allowed the functional comparison of the delimited potential NT locus.

The genomic region identified by MBS was further restricted by designing genetic markers that do not show recombination with the phenotype on extreme F2 genotypes analyzed together with susceptible and tolerant standard lines (Additional file S2D). This interval spans regions of 0.7 Mbp and 0.76 Mbp in U2BvONT and 2320BvONT, respectively. Since the genome sequence is continuous and colinear in both assemblies at the potential NT locus, there is no evidence for missing additional sequences.

Genome sequences and DNA sequence data in general enable sequence comparison to transfer gene function information from one species to another. For example, several *Arabidopsis thaliana* genes were found to be homologous to the *Lotus japonicus* gene *NODULE INCEPTION* [22] encoding the plant-specific transcription factor (TF) LjNIN [23]. The *A. thaliana* NIN-LIKE PROTEIN (NLP) family of TFs control nitrate-responsive gene transcription [24]. Among the NLPs is AtNLP7 (At4g24020) which is well described as a major regulator of nitrate signaling [25–29]. Evidence for direct target genes regulated by AtNLP7 has recently been published [25]. NLPs including AtNLP7 contain the sequence-conserved PB1 domain (NCBI domain cd06407; [30]) which mediates protein-protein interactions [31]. These include NLP-NLP interactions as well as interactions between NLPs and other factors [29,32]. Four core amino acid residues within the PB1 domain (K867, D909, D911, and E913) are thought to be required for NLP-NLP interactions [32]. In the context of nitrate response and plant growth, mutants with substitutions of these core amino acid residues require a higher level of expression than wildtype NLP7 [32]. Another highly conserved residue in NLPs is S205 which serves as regulatory phosphorylation site [33].

The potential NT locus shows over the length of about of 0.7 Mbp a generally high synteny between the genome assemblies U2BvONT and 2320BvONT. Within this 0.7 Mbp region, we identified a cluster of genes related to *AtNLP7* that spans less than 100 kbp (Figure 3). Within the whole genome sequence, *BvNLP7* genes were only found at the potential NT locus on chr5. In the published *ab initio* structural annotation of EL10, two additional gene models (compared to the liftoff from the U2BvONT annotation, Figure 3) were predicted. However, both show similarity to histone H4 and are not considered as candidates.

Despite the synteny within the target region, four *BvNLP7* genes were identified in U2BvONT, but only three in 2320BvONT. The core amino acid residues of the NLP7 PB1 domain, which is relevant for regulatory interactions, are conserved in almost all of the *BvNLP7* genes.

Nitrogen is not only a major nutrient for plants but is also needed as signaling molecule for developmental processes and defense against pathogens [27,34]. For example, the signaling molecule nitric oxide mediates defense responses and nitric oxide production is therefore increased during plant-pathogen interaction [35]. In previous studies, reduced nitrate uptake and transport capacity was observed upon nematode infection of *Coffea arabica* plants and might be the direct result of root damage caused by the activities (including feeding) of the nematodes [34,36]. As the plant has to adapt to the changing conditions caused by the nematodes, which deprive the plant of nitrogen, adaptation of the plants nitrogen metabolism is crucial [34]. Indeed, NLP7 is known to be involved in the control of plant nitrate metabolism [27]. A high number of genes involved in nitrate signaling are among the targets of NLP7. As nitrate signaling is disturbed upon nematode infection, NLP7 might be able to modulate gene expression allowing the plant to adapt and to better cope with the infection, which fits to the observed tolerance.

In the context of nitrate response and plant growth, mutants with substitutions of the core amino acid residues described above require a higher level of expression than wildtype NLP7 [32]. The fact that *nlp7* mutants are associated with damaged roots as well as impaired nitrate assimilation leading to decreased amino acid formation [26] might at least partially explain the trade-off between high yield and resistance against nematodes.

To get insights into expression patterns of the *BvNLP7* genes and to get an overview of the response of resistant roots to the infection with *H. schachtii*, an infection assay was performed. RNA-Seq analysis was used to identify up-and downregulated genes in response to BCN infection (21 dpi). Recently, a detailed overview over the transcriptional responses during the beet-nematode interaction has been published [6], addressing nematode effector genes related to tolerance and resistance. In the resistant cultivar ’Nemakill’, an induction of genes related to plant defense response was observed. For all DEGs for the comparisons i) SUS3 treated vs BR12 treated, ii) all tolerant vs all susceptible, and iii) U1Bv treated vs U2Bv treated, we looked at the gene’s functional annotation (Additional file S1G, Additional file S1H, Additional file S1I) and compared these functions to the findings of Ghaemi *et al.* [6]. Gene expression differences in phytohormone-related genes as well as genes involved in the plant defense response and the phenylpropanoid pathway were reported between Nemakill and a susceptible cultivar. Such genes are also frequently present among the DEGs we detected in our study for all three comparisons (see above). Also, CYSTM domain-encoding genes were found to be upregulated in infected Nemakill roots [6], but such genes were not detected in our study. Chitinases, F-box proteins and CASP-like proteins have been reported to be involved in the defense mechanism against nematodes, in various processes including plant-pathogen interactions, and in stress resistance against nematodes, respectively. Genes encoding these types of proteins have been found in both RNA-Seq based studies, the one using Nemakill [6] as well as in our study for the three comparisons. Overall, there is a significant overlap between the potentially relevant genes reported by in Ghaemi *et al.* [6] and the DEGs detected in our study. This suggests that a broad and strong defense response is triggered in the resistant roots upon infection with nematodes.

A time course experiment might reveal additional insights into the plants’ defense mechanisms upon infection. However, the performed experiment enabled an in depth, transcriptomic characterization of our *BvNLP7* genes. All four U2BvONT *BvNLP7* genes are expressed. Expression is significantly higher in inoculated tolerant BR12 samples compared to inoculated susceptible SUS3 standard samples. The genes *NLP7-T1* and *NLP7-T2* are significantly differentially expressed between all samples of tolerant lines and all samples of susceptible lines.

Comparison with other nematode-*A. thaliana* transcriptomic datasets using the tool NEMATIC [37] revealed that *AtNLP7* (At4g24020) is downregulated in syncytia [4] (at 5 dpi and 15 dpi Syncytium vs Root, see Table S1) and non-infected root samples. Further, the analyses showed that *AtNLP7* is expressed in different parts of the roots, for example in the root hair zone and in lateral roots.

Genes directly regulated by the AtNLP7 TF in *A. thaliana* root cells [25] include several genes encoding transporters, expansins and a WRKY TF (WRKY23) which are involved in nematode-induced syncytia formation. On chr5 of U2BvONT and 2320BvONT, 12 genes were annotated as putative WRKY TFs. Two (Bv05_g11386_pnrz and Bv05_g11531_wmfg) are located close to the potential NT locus at approximately 1 Mbp up-and downstream of the *BvNLP7* cluster, respectively. Two additional WRKY23 homologs (Bv05_g14901_ygdi and Bv05_g14902_sogd) were identified on chr5. However, none of these four WRKYs are differentially expressed in the comparisons of all tolerant vs. all susceptible samples or inoculated tolerant line BR12 vs. inoculated susceptible line SUS3.

In oilseed rape, BCN resistance was enhanced by gene pyramiding [38]. Homologs of the plant-defense genes addressed by Zhong *et al*., namely *AtNPR1/AT1G64280*, *AtSGT1b/AT4G11260* and *AtRAR1/AT5G51700*, were identified in U2BvONT. BvNPR1 was detected to be encoded on chr8 with 58.6% aa identity. Homologs of the other two genes were found to be encoded on chr3 with 57.7% (*BvSGT1b*) and 63.6% (*BvRAR1*) sequence identity at codon/aa level. In another recent study, BCN infection phenotypes were characterized in transgenic *A. thaliana* [39]. A *AtSNAP2*/*AT3G56190* homolog was detected to be encoded on U2BvONT chr2 (74.9% identity via blastp), a *AtSHM(T)4*/*AT4G13930* homolog on chr3 (89.2% aa identity) and a *AtPR1/AT2G14610* homolog on chr9 (58.6% aa identity). In summary, none of the sugar beet homologs of these six genes which were described to be associated with BCN infection/resistance in these two studies, are located within or next to the potential NT locus. In addition, these genes are not differentially expressed when comparing all tolerant vs. all susceptible samples. Therefore, these genes are highly unlikely to be causal for BCN tolerance/resistance.

In summary, the presence of one more *BvNLP7* (additional) gene in Strube U2Bv as well as the cumulative expression of those genes might explain the BCN tolerance of the genotype Strube U2Bv. Therefore, the BCN tolerance might be based on a ‘gene dosage’ effect.

## 4 Conclusions

In this study, a potential trait locus associated with BCN tolerance was identified via mapping-by-sequencing. Two newly generated long read-based genome sequences of the sugar beet reference genotype 2320Bv and the tolerant line Strube U2Bv guided the characterization of the potential NT locus. Four *BvNLP7* genes in U2BvONT are upregulated in tolerant lines as revealed by an infection assay. These four genes, *NLP7-T1-4*, combined, or a subset thereof, might convey tolerance against the cyst nematode *H. schachtii* which infects sugar beet and is a serious problem due to yield loss. These results have positive implications for knowledge-based breeding of elite genotypes.

## 5 Methods

### 5.1 Plant material and growth conditions

The breeding material is a large population of 406 lines descending from a single F1 plant obtained by crossing a highly BCN susceptible line (Strube U1Bv) with a line (Strube U2BV) highly tolerant to *H. schachtii*. Leaf samples were collected from all F2 plants and frozen at − 20 °C until usage. Ten selfed F3 plants per line were grown at a 16 hours light, 8 hours dark cycle at 20-22 °C in the greenhouse, and 4,060 plants were used for evaluating nematode tolerance. Single seedlings were grown in folding boxes (40×20×120 mm) that guarantee separation of the root system and all plants were randomized in larger boxes (41×26×13 cm) accommodating 120 plants each together with tolerant and susceptible *H. schachtii* breeding material used as checks. Four months after sowing, each plant was inoculated with 350 *H. schachtii* second stage juvenile larvae using a dispenser, following an internal protocol of Strube Research. Five weeks after inoculation, roots of each plant were washed, and cysts were collected and counted under a stereoscope.

For RNA-Seq, different BCN tolerant and susceptible lines were used. The susceptible lines were Strube U1Bv and SUS3, an internal standard line of Strube Research. The tolerant lines were Strube U2Bv and the best performing F2 genotype BR12. All lines were germinated and cultivated in the greenhouse for 11 weeks in total. Each line was represented by 40 plants. Half of the plants were inoculated with nematodes as described above, the other half was left untreated. Inoculation was done after 8 weeks, and all material was collected at 21 dpi (3 weeks after inoculation). The sampling of infected plants was performed by collecting tissue and washing off the cysts, which were then counted under a stereomicroscope. After gentle washing, the shoot was removed and the roots were immediately frozen in liquid nitrogen until RNA extraction.

The phenotypic data were analyzed using a mixed model approach [40]. Boxes were treated as random incomplete blocks and the genotypes as fixed. The counted number of cysts (n) was transformed using a square root transformation (SN = Jn + 3/8) to meet the assumption of normally distributed residuals required for mixed models. Adjusted means were obtained for each line for QTL mapping. Adjusted single plant data were used to explore within-family variation using the CV. A small CV indicates phenotypic homogeneity among individual plants of a line, suggesting no segregation for the trait under study.

### 5.2 QTL detection

Together with the genetic linkage map, the adjusted means of the F2:3 families of the STR-NT population were used for QTL mapping. Composite interval mapping (CIM) was employed for QTL detection and a LOD threshold of 3.5 corresponding to an experiment wise type I error rate of 0.05 was determined using 1000 permutation runs. All QTL computations were performed with the software package PLABQTL [41] using an additive and dominant model and a scan of 1 cM interval.

### 5.3 DNA extraction and MBS pool generation

Genomic DNA for MBS was extracted from young leaf tissue using the CTAB method [42]. The nine most susceptible genotypes were used for individual library preparations, whereas the gDNA from the remaining seven genotypes were equimolarly pooled before library preparation (16 lines in total). Of these and with regard to SNP192, 8 carry only the C allele (susceptible), 1 carries only the G allele (tolerant) and 7 were heterozygous. For the ‘tolerant’ pool, the gDNA from the nine most tolerant lines was used for individual library preparations, whereas the remaining 11 gDNAs were equimolarly pooled (20 lines in total). Of these and with regard to SNP192, 14 carry only the G allele (tolerant), 1 carries only the C allele (susceptible) and 5 were heterozygous. DNA for short read sequencing and PCR-based marker analysis was extracted from 8 leaf disks with 1 cm diameters using a CTAB-based protocol [42]. High molecular weight DNA for long read ONT sequencing was extracted with a modified CTAB-based protocol as previously described [43].

### 5.4 Short read sequencing for MBS

Each single gDNA or gDNA pool was fragmented by sonication using a Bioruptor (Fa. Diagenode). After cleaning the DNAs by AMPureXP Beads (Fa. Beckmann-Coulther), 200 ng of fragmented DNA was used for library preparation with the TruSeq Nano DNA library preparation kit (Fa. Illumina). End-repaired fragments were size selected by AMPureXP Beads to an average size of around 700 bp. After A-tailing and ligation of barcoded adaptors, fragments were enriched by 8 cycles of PCR. The final libraries were quantified using PicoGreen (Fa. Quant-iT) on a FLUOstar plate reader (Fa. BMG labtech) and quality checked by HS-Chip on a 2100 Bioanalyzer (Fa. Agilent Technologies). After pooling of all libraries, sequencing was performed on two 2×250 nt runs on a HiSeq1500 in rapid mode over two lanes using onboard-cluster generation. Processing and demultiplexing of raw data were performed by bcl2fastq. Additional file S2G summarizes the data submitted to ENA.

### 5.5 Short read mapping and variant calling from MBS data

The short read WGS data from the phenotypic pools, the parental lines of the mapping population, the F1 plant, and additional standard lines were used to identify small sequence variations within the population and against the susceptible reference genotype 2320Bv. BWA MEM v0.7.13 [44] was applied with the –m option to flag small alignments as secondary to align short reads to the U2BvONT reference sequence. Picard tools v2.5.0 (https://broadinstitute.github.io/picard/) and samtools v1.15.1 [45] were used to mark PCR duplicates, sort, and index BAM files. Mappings were filtered with samtools to remove spurious hits, low quality alignments, and reads that are not properly mapped in pairs (-q 30 -b -F 0×900 -f 0×2). GATK v3.8 [46] [47] was applied for the detection of small sequence variants as previously described [48]. Sequence variants were filtered to obtain a reduced set with high confidence. The following criteria were applied to select high confidence variants: (1) variants are homozygous in the parent reads, (2) variants are contrasting between the parents, and (3) variants are heterozygous in the F1 reads. Python scripts developed and applied for filtering are available in the corresponding GitHub repository (https://github.com/bpucker/beetresmabs).

### 5.6 Calculation of delta allele frequencies and interval detection

The calculation and analysis of dAFs is generally based on a previously described workflow [49]. The methods are described in detail in Additional file S2H. Figure 3, which represents the comparison of the potential NT locus, was mostly generated with gggenomes [50].

### 5.7 RNA Extraction, Library Preparation, and Sequencing

Plants of the genotypes Strube U2Bv, BR12, Strube U1Bv and SUS3 were grown in the greenhouse and either infected or not infected with *H. schachtii* as described in 5.1. In total, 24 samples (Additional file S2I) including three biological replicates for each condition of the infection experiment were ground separately under liquid nitrogen. Total RNA was extracted from approx. 100 mg tissue using an RNA Isolation Kit (Sigma-Aldrich Spectrum™ Plant Total RNA) according to suppliers’ instructions. Quality control, determination of RIN numbers, and estimation of the concentrations of RNA samples was done on a Bioanalyzer 2100 (Agilent) using RNA Nano 6000 Chips. To construct sequencing libraries according to the Illumina TruSeq RNA Sample Preparation v2 Guide, 500 ng total RNA per subsample were used. Further steps, like enrichment of poly-A containing mRNA, cDNA synthesis, adapter ligation, PCR enrichment, library quantification, and equimolar pooling, were performed according to Theine *et al.*, 2021 [51]. Single end sequencing of 100 nt was performed on an Illumina NextSeq500.

### 5.8 Genetic markers and linkage mapping

Sets of small variants detected by GATK v3.8 [46,47], on the basis of short read mappings of Strube U1Bv, Strube U2Bv, and F1 reads to RefBeet-1.2 [16], were used to design KASPar markers. Only homozygous single nucleotide variants contrasting between the parents and with clear heterozygosity in the F1 were taken forward as clean variants. All variants overlapping with existing markers were excluded. Up to 1000 variants per contig were selected based on the quality of the variant call. A total of 50 bp upstream and downstream, respectively, were checked for other variants based on a very lenient and unfiltered variant calling. Only marker candidates without any additional variants in these flanking sequences were taken forward. Further, only marker candidates with less than 65% GC content in 100 bp of flanking sequence were considered. Finally, marker candidates were preferentially selected on unplaced contigs of RefBeet-1.2 with an upper limit of four candidates per contig. The markers included within the final list were selected based on their position on each chromosome aiming to form a well-distributed marker subset.

The linkage map was constructed using KASPar markers and the package R/qtl [52]. The final linkage map comprises a set of 194 SNP markers including SNP192 that coalesced into nine linkage groups. Each group corresponded to one of the nine chromosomes in the haploid sugar beet genome. The average distance between loci was 3 cM except for two markers at the end of chr1 in poor linkage due to distortion. The average number of markers per chromosome was 21. For chr7 and chr9 the number of markers was below average and equal to 15 and 11, respectively.

### 5.9 ONT sequencing and ONT assembly

ONT long-read sequencing was performed on a GridION. The initial assembly was generated with Canu [53] and further processed. The exact methodology is described in Additional file S2H, and Additional file S2G summarizes the data submitted to ENA.

### 5.10 Gene prediction and functional annotation

After repeat masking, hint-based gene prediction was performed mainly with BRAKER2 [54]. All predicted genes were subsequently functionally annotated. The methodology for structural and functional annotation with BRAKER2 is described in detail in Additional file S2H.

### 5.11 Differential gene expression analysis

The generated RNA-Seq reads were mapped to the U2BvONT genome sequence using STAR v2.7.6a (Linux_x86_64_static/STARlong --runThreadN 8 --genomeDir /dir/ -- limitBAMsortRAM 32000000000 --outBAMsortingThreadN 4 --outSAMtype BAM SortedByCoordinate --outFileNamePrefix /01 --readFilesCommand gunzip -c –readFilesIn 01.fastq.gz) [55]. FeatureCounts v2.0.0 [56] was used to quantify annotated genes in the U2BvONT GFF file (-T 8 -t gene -a annotation.gff -o readcounts_allbams.txt *.bam). Downstream analysis was performed using the R package DESeq2 v1.26.0 [57]. A variance stabilizing transformation was conducted. A principal component analysis (PCA) for all samples of the infection assay (Additional file S2F) was generated using prcomp (stats-package v3.6.3 [58] and ggplot2 v3.3.5 [59]. Significantly differentially expressed genes between i) all tolerant vs. all susceptible samples and ii) between inoculated SUS3 and inoculated BR12 samples, were extracted based on an adjusted p-value < 0.05.

### 5.12 Rearrangement and synteny analyses

Synteny analyses between 2320BvONT and U2BvONT as well as 2320BvONT and RefBeet were performed using JCVI MCscan v1.2.4 [60]. Unanchored mRNAs were compared for unique functions using BLAST v2.13.0 [61] and InterProScan v5.52 [62]. Structural rearrangements were identified with SyRI v1.4 [63].

### 5.13 Comparison of *BvNLP7* genes

A list of direct targets of the AtNLP7 TF has been published recently [25]. To assess a possible role of NPL7 gene(s) in BCN tolerance, these targets were functionally investigated by overrepresentation analysis of GO terms using PANTHER v17.0 [64]. Additionally, the sequences were directly compared via MAFFT v7.487 [65] alignments and manual inspection.

## 6 Declarations

### Ethics approval and consent to participate

All sugar beet material relevant for the BCN studies was provided by Strube Research GmbH & Co. KG, KWS2320 material was provided by KWS Saat ES. The material of Strube and KWS was transferred under the regulations of a standard material transfer agreement (SMTA) according to the International Treaty. Plants were grown in accordance with German legislation.

### Consent for publication

Not applicable.

### Availability of data and materials

A summary of the availability of the newly generated sequencing data, assemblies and annotations is provided in Additional file S2G covering ENA projects PRJEB56338, PRJEB37059, PRJEB36905, PRJEB58360 as well as DOIs for the genome sequence annotation files (.gff). Additional RNA-Seq datasets produced for hint generation and gene prediction are available via ENA projects PRJEB58621 and PRJEB62793, these and further already published hint data are listed in Additional file S2J.

## Funding

This project was funded by the Federal Ministry of Education and Research of Germany (BMBF) in the frame of KMU-innovativ-16 under the grant numbers 031B0081A and 031B0081B (BeetRes-MaBS). This work was supported by the BMBF-funded de.NBI Cloud within the German Network for Bioinformatics Infrastructure (grant numbers 031A532B, 031A533A, 031A533B, 031A534A, 031A535A, 031A537A, 031A537B, 031A537C, 031A537D, 031A538A). KS is funded by Bielefeld University through the Graduate School DILS (Digital Infrastructure for the Life Sciences). We acknowledge support for the publication costs by the Deutsche Forschungsgemeinschaft and the Open Access Publication Fund of Bielefeld University.

### Authors’ contributions

EO, AM, AS, DH and BW are responsible for conceptualization. KS, BP, EO, EA, AS, DH and BW developed the methodology. Validation, formal analysis and investigation were performed by KS, BP, EO, LS, PV, EA and DH. KS, BP, EO, LS, PV, EA and DH conducted data curation. KS, BP, EO, and DH wrote the original draft. KS, BP, EO, EA, AM, AS, PV, BW and DH reviewed and edited the manuscript. Visualization of the data was performed by KS and BP. Supervision and project administration was carried out by EO, AS, DH and BW. EO, AM, AS, and BW were responsible for funding acquisition.

## Acknowledgements

We thank the CeBiTec Bioinformatic Resource Facility team as well as the de.NBI team for great technical support. The authors also wish to thank the members of Strube Research for their technical support in phenotyping and genotyping large populations. We thank Prof. Piepho from the University of Hohenheim for the constructive discussion on the liner mixed model. We thank Britta Schulz from KWS-SAAT SE for providing seeds of the reference genotype KWS2320. Thanks to Julia Zimmer and Marvin Hildebrandt for variant validation.

## 7 Supplementary Material

**Additional file S1: S1A:** Genomic positions of the new set of 187 markers designed based on data for Strube U2Bv. In the first column, the name of the marker is shown, followed by the chromosome and genomic position for each assembly (2320BvONT v1.0 and U2BvONT v1.0). **S1B:** Overrepresentation analysis (GO terms) of the AtNLP7 targets identified by Alvarez et al., 2020. **S1C:** Percent identity matrices for the BvNLP7 genes. **S1D:** List of all significantly differentially expressed genes for the comparison all tolerant vs all susceptible. **S1E:** List of all significantly differentially expressed genes for the comparison SUS3 treated vs BR12 treated. **S1F:** List of all significantly differentially expressed genes for the comparison U1Bv treated vs U2Bv treated. **S1G:** List of functionally annotated DEGs for the comparison all tolerant vs all susceptible. **S1H:** List of functionally annotated DEGs for the comparison SUS3 treated vs BR12 treated. **S1I:** List of functionally annotated DEGs for the comparison U1Bv treated vs U2Bv treated.

**Additional file S2: S2A:** Histogram of adjusted cyst counts (SN) data per family. **S2B:** Structural rearrangements between 2320BvONT and U2BvONT as identified with SyRI. **S2C:** Delta allele frequency plots for 10 SNP windows of all nine U2BvONT pseudochromosomes. **S2D:** Marker information/primer sequences used for delimitation of the potential NT locus and graphical genotyping with flanking and co-segregating markers. **S2E:** Visualization of key polymorphisms at conserved positions of Arabidopsis NLP7. **S2F:** Principal component analysis for all samples of the RNA-Seq infection experiment. **S2G:** Availability and composition of datasets generated. **S2H:** Detailed methods for i) calculation of the delta allele frequencies and interval detection, ii) ONT sequencing, iii) ONT assembly, and iv) gene prediction and functional annotation. **S2I:** Overview of the RNA-Seq samples in the infection experiment. **S2J:** List of all RNA-Seq datasets incorporated as hints for the gene prediction including two newly submitted datasets. **S2K:** Dot plot heatmap of the closed gap region in the initial Strube U2Bv assembly.

**Additional file S3:** Multiple sequence alignment of coding sequences of all *BvNLP7* genes.

**Additional file S4:** Multiple sequence alignment of *A. thaliana* NLP7 and U2BvONT and 2320BvONT rycp aa sequences.

**Additional file S5:** Multiple sequence alignment of aa sequences encoded by the *BvNLP7* genes (without rycp).

**Additional file S6:** Multiple sequence alignment of aa sequences encoded by all BvNLP7 genes.

## References

1. Fischer, H.E. Origin of the “Weisse Schlesische Ruebe” (white Silesian beet) and resynthesis of sugar beet. Euphytica 41, 75–80 10.1007/BF00022414 (1989).

2. Biancardi, E., McGrath, J.M., Panella, L.W., Lewellen, R.T. & Stevanato, P. Sugar Beet in Root and Tuber Crops Vol. 7 (ed Bradshaw, J.E.) 173-219 Ch. Chapter 6 10.1007/978-0-387-92765-7_6 (Springer, 2010).

3. Wieczorek, K., Golecki, B., Gerdes, L., Heinen, P., Szakasits, D., Durachko, D.M., Cosgrove, D.J., Kreil, D.P., Puzio, P.S., Bohlmann, H. & Grundler, F.M. Expansins are involved in the formation of nematode-induced syncytia in roots of Arabidopsis thaliana. The Plant Journal 48, 98–112 10.1111/j.1365-313X.2006.02856.x (2006).

4. Szakasits, D., Heinen, P., Wieczorek, K., Hofmann, J., Wagner, F., Kreil, D.P., Sykacek, P., Grundler, F.M. & Bohlmann, H. The transcriptome of syncytia induced by the cyst nematode Heterodera schachtii in Arabidopsis roots. *The Plant Journal* **57**, 771-84 10.1111/j.1365-313X.2008.03727.x (2009).

5. Wieczorek, K., Hofmann, J., Blochl, A., Szakasits, D., Bohlmann, H. & Grundler, F.M. Arabidopsis endo-1,4-beta-glucanases are involved in the formation of root syncytia induced by Heterodera schachtii. *The Plant Journal* **53**, 336-51 10.1111/j.1365-313X.2007.03340.x (2008).

6. Ghaemi, R., Pourjam, E., Safaie, N., Verstraeten, B., Mahmoudi, S.B., Mehrabi, R., De Meyer, T. & Kyndt, T. Molecular insights into the compatible and incompatible interactions between sugar beet and the beet cyst nematode. BMC Plant Biology 20, 483 10.1186/s12870-020-02706-8 (2020).

7. Richardson, K.L. Registration of sugar beet mapping populations CN239, CN240, and CN241 segregating for resistance to Heterodera schachtii from sea beet. *Journal of Plant Registrations* 16, 459-464 10.1002/plr2.20152 (2022).

8. Vandenbossche, B.A.B., Niere, B. & Vidal, S. Effect of Temperature on the Hatch of Two German Populations of the Beet Cyst nematodes, Heterodera schachtii and Heterodera betae. Journal of Plant Diseases and Protection 122, 250–254 10.1007/BF03356560 (2016).

9. Cai, D., Kleine, M., Kifle, S., Harloff, H.J., Sandal, N.N., Marcker, K.A., Klein Lankhorst, R.M., Salentijn, E.M.J., Lange, W., Stiekema, W.J., Wyss, U., Grundler, F.M.W. & Jung, C. Positional cloning of a gene for nematode resistance in sugar beet. *Science* **275**, 832-834 10.1126/science.275.5301.832 (1997).

10. Cai, D., Thurau, T., Tian, Y., Lange, T., Yeh, K.W. & Jung, C. Sporamin-mediated resistance to beet cyst nematodes (Heterodera schachtii Schm.) is dependent on trypsin inhibitory activity in sugar beet (Beta vulgaris L.) hairy roots. Plant Molecular Biology 51, 839–49 10.1023/a:1023089017906 (2003).

11. Reuther, M., Lang, C. & Grundler, F.M.W. Nematode-tolerant sugar beet varieties – resistant or susceptible to the Beet Cyst Nematode Heterodera schachtii? Sugar Industry 142, 277–284 10.36961/si18397 (2017).

12. Holtmann, B., Kleine, M. & Grundler, F.M.W. Ultrastructure and anatomy of nematode-induced syncytia in roots of susceptible and resistant sugar beet. Protoplasma 211, 39–50 10.1007/BF01279898 (2000).

13. Stevanato, P., Trebbi, D., Panella, L., Richardson, K., Broccanello, C., Pakish, L., Fenwick, A.L. & Saccomani, M. Identification and Validation of a SNP Marker Linked to the Gene HsBvm-1 for Nematode Resistance in Sugar Beet. Plant Molecular Biology Reporter 33, 474–479 10.1007/s11105-014-0763-8 (2015).

14. Biancardi, E., Lewellen, R.T., Frese, L., Ford-Lloyd, B., de Biaggi, M., Hautekeete, N., van Dijk, H., Touzet, P., Bartsch, D., Panella, L.W., Stevanato, P., Pavli, O., Skaracis, G. & McGrath, J.M. Beta maritima: The Origin of Beets (Second Edition) 10.1007/978-3-030-28748-1 2020).

15. McGrath, J.M., Funk, A., Galewski, P., Ou, S., Townsend, B., Davenport, K., Daligault, H., Johnson, S., Lee, J., Hastie, A., Darracq, A., Willems, G., Barnes, S., Liachko, I., Sullivan, S., Koren, S., Phillippy, A., Wang, J., Liu, T., Pulman, J., Childs, K., Shu, S., Yocum, A., Fermin, D., Mutasa-Gottgens, E., Stevanato, P., Taguchi, K., Naegele, R. & Dorn, K.M. A contiguous de novo genome assembly of sugar beet EL10 (Beta vulgaris L.). DNA Research 30, 10.1093/dnares/dsac033 (2023).

16. Dohm, J.C., Minoche, A.E., Holtgrawe, D., Capella-Gutierrez, S., Zakrzewski, F., Tafer, H., Rupp, O., Sorensen, T.R., Stracke, R., Reinhardt, R., Goesmann, A., Kraft, T., Schulz, B., Stadler, P.F., Schmidt, T., Gabaldon, T., Lehrach, H., Weisshaar, B. & Himmelbauer, H. The genome of the recently domesticated crop plant sugar beet (Beta vulgaris). Nature 505, 546–9 10.1038/nature12817 (2014).

17. Holtgräwe, D., Sörensen, T.R., Viehöver, P., Schneider, J., Schulz, B., Borchardt, D., Kraft, T., Himmelbauer, H. & Weisshaar, B. Reliable in silico identification of sequence polymorphisms and their application for extending the genetic map of sugar beet (Beta vulgaris). *PLoS ONE* **9**, e110113 10.1371/journal.pone.0110113 (2014).

18. Rodriguez Del Rio, A., Minoche, A.E., Zwickl, N.F., Friedrich, A., Liedtke, S., Schmidt, T., Himmelbauer, H. & Dohm, J.C. Genomes of the wild beets Beta patula and Beta vulgaris ssp. maritima. *The Plant Journal* **99**, 1242-1253 10.1111/tpj.14413 (2019).

19. Smit, A.F.A., Hubley, R. & Green, P. RepeatMasker Open-4.0. (2015).

20. Ries, D., Holtgräwe, D., Viehöver, P. & Weisshaar, B. Rapid gene identification in sugar beet using deep sequencing of DNA from phenotypic pools selected from breeding panels. *BMC Genomics* 17, 236 10.1186/s12864-016-2566-9 (2016).

21. Flavell, R.B., Bennett, M.D., Smith, J.B. & Smith, D.B. Genome size and the proportion of repeated nucleotide sequence DNA in plants. Biochemical Genetics 12, 257–69 10.1007/BF00485947 (1974).

22. Schauser, L., Roussis, A., Stiller, J. & Stougaard, J. A plant regulator controlling development of symbiotic root nodules. Nature 402, 191–5. 10.1038/46058 (1999).

23. Schauser, L., Wieloch, W. & Stougaard, J. Evolution of NIN-like proteins in Arabidopsis, rice, and Lotus japonicus. Journal of Molecular Evolution 60, 229–37 10.1007/s00239-004-0144-2 (2005).

24. Konishi, M. & Yanagisawa, S. Arabidopsis NIN-like transcription factors have a central role in nitrate signalling. *Nature Communications* 4, 1617 10.1038/ncomms2621 (2013).

25. Alvarez, J.M., Schinke, A.L., Brooks, M.D., Pasquino, A., Leonelli, L., Varala, K., Safi, A., Krouk, G., Krapp, A. & Coruzzi, G.M. Transient genome-wide interactions of the master transcription factor NLP7 initiate a rapid nitrogen-response cascade. *Nature Communications* **11**, 1157 10.1038/s41467-020-14979-6 (2020).

26. Castaings, L., Camargo, A., Pocholle, D., Gaudon, V., Texier, Y., Boutet-Mercey, S., Taconnat, L., Renou, J.P., Daniel-Vedele, F., Fernandez, E., Meyer, C. & Krapp, A. The nodule inception-like protein 7 modulates nitrate sensing and metabolism in Arabidopsis. The Plant Journal 57, 426–35 10.1111/j.1365-313X.2008.03695.x (2009).

27. Marchive, C., Roudier, F., Castaings, L., Brehaut, V., Blondet, E., Colot, V., Meyer, C. & Krapp, A. Nuclear retention of the transcription factor NLP7 orchestrates the early response to nitrate in plants. *Nature Communications* **4**, 1713 10.1038/ncomms2650 (2013).

28. Zhao, L., Zhang, W., Yang, Y., Li, Z., Li, N., Qi, S., Crawford, N.M. & Wang, Y. The Arabidopsis NLP7 gene regulates nitrate signaling via NRT1.1-dependent pathway in the presence of ammonium. *Scientific Reports* **8**, 1487 10.1038/s41598-018-20038-4 (2018).

29. Guan, P., Ripoll, J.J., Wang, R., Vuong, L., Bailey-Steinitz, L.J., Ye, D. & Crawford, N.M. Interacting TCP and NLP transcription factors control plant responses to nitrate availability. *Proceedings of the National Academy of Sciences of the United States of America* **114**, 2419-2424 10.1073/pnas.1615676114 (2017).

30. Lu, S., Wang, J., Chitsaz, F., Derbyshire, M.K., Geer, R.C., Gonzales, N.R., Gwadz, M., Hurwitz, D.I., Marchler, G.H., Song, J.S., Thanki, N., Yamashita, R.A., Yang, M., Zhang, D., Zheng, C., Lanczycki, C.J. & Marchler-Bauer, A. CDD/SPARCLE: the conserved domain database in 2020. Nucleic Acids Research 48, D265–D268 10.1093/nar/gkz991 (2020).

31. Sumimoto, H., Kamakura, S. & Ito, T. Structure and function of the PB1 domain, a protein interaction module conserved in animals, fungi, amoebas, and plants. SCIENCE’S STKE Science Signaling 2007, re6 10.1126/stke.4012007re6 (2007).

32. Konishi, M. & Yanagisawa, S. The role of protein-protein interactions mediated by the PB1 domain of NLP transcription factors in nitrate-inducible gene expression. BMC Plant Biology 19, 90 10.1186/s12870-019-1692-3 (2019).

33. Liu, K.H., Niu, Y., Konishi, M., Wu, Y., Du, H., Sun Chung, H., Li, L., Boudsocq, M., McCormack, M., Maekawa, S., Ishida, T., Zhang, C., Shokat, K., Yanagisawa, S. & Sheen, J. Discovery of nitrate-CPK-NLP signalling in central nutrient-growth networks. Nature 545, 311–316 10.1038/nature22077 (2017).

34. Labudda, M., Różańska, E., Muszyńska, E., Marecka, D., Głowienka, M., Roliński, P. & Prabucka, B. Heterodera schachtii infection affects nitrogen metabolism in Arabidopsis thaliana. *Plant Pathology* **69**, 794-803 10.1111/ppa.13152 (2020).

35. Salgado, I., Modolo, L.V., Augusto, O., Braga, M.R. & Oliveira, H.C. Mitochondrial Nitric Oxide Synthesis During Plant–Pathogen Interactions: Role of Nitrate Reductase in Providing Substrates in Nitric Oxide in Plant Growth, Development and Stress Physiology (eds Lamattina, L. & Polacco, J.C.) 239-254 Ch. Chapter 95 10.1007/7089_2006_095 2006).

36. Vaast, P., Caswell-Chen, E.P. & Zasoski, R.J. Effects of two endoparasitic nematodes (Pratylenchus coffeae and Meloidogyne konaensis) on ammonium and nitrate uptake by Arabica coffee (Coffea arabica L.). Applied Soil Ecology 10, 171–178 10.1016/S0929-1393(98)00037-7 (1998).

37. Cabrera, J., Bustos, R., Favery, B., Fenoll, C. & Escobar, C. NEMATIC: a simple and versatile tool for the in silico analysis of plant-nematode interactions. *Molecular Plant Pathology* 15, 627-36 10.1111/mpp.12114 (2014).

38. Zhong, X., Zhou, Q., Cui, N., Cai, D. & Tang, G. BvcZR3 and BvHs1(pro-1) Genes Pyramiding Enhanced Beet Cyst Nematode (Heterodera schachtii Schm.) Resistance in Oilseed Rape (Brassica napus L.). International Journal of Molecular Sciences 20, 10.3390/ijms20071740 (2019).

39. Zhao, J., Duan, Y., Kong, L., Huang, W., Peng, D. & Liu, S. Opposite Beet Cyst Nematode Infection Phenotypes of Transgenic Arabidopsis Between Overexpressing GmSNAP18 and AtSNAP2 and Between Overexpressing GmSHMT08 and AtSHMT4. *Phytopathology* **112**, 2383-2390 10.1094/PHYTO-01-22-0011-R (2022).

40. Piepho, H.P., Buchse, A. & Emrich, K. A Hitchhiker’s Guide to Mixed Models for Randomized Experiments. *Journal of Agronomy and Crop Science* **189**, 310-322 10.1046/j.1439-037X.2003.00049.x (2003).

41. Utz, H.F. & Melchinger, A.E. PLABQTL: a program for composite interval mapping of QTL. Journal of Agricultural Genomics 2, 1–6 (1996).

42. Rosso, M.G., Li, Y., Strizhov, N., Reiss, B., Dekker, K. & Weisshaar, B. An *Arabidopsis thaliana* T-DNA mutagenised population (GABI-Kat) for flanking sequence tag based reverse genetics. *Plant Molecular Biology* **53**, 247-259 10.1023/B:PLAN.0000009297.37235.4a (2003).

43. Siadjeu, C., Pucker, B., Viehöver, P., Albach, D.C. & Weisshaar, B. High Contiguity De Novo Genome Sequence Assembly of Trifoliate Yam (Dioscorea dumetorum) Using Long Read Sequencing. Genes 11, E274 10.3390/genes11030274 (2020).

44. Li, H. Aligning sequence reads, clone sequences and assembly contigs with BWA-MEM. arXiv https://arxiv.org/abs/1303.3997, posted 2013-05-26 10.48550/arXiv.1303.3997 (2013).

45. Li, H., Handsaker, B., Wysoker, A., Fennell, T., Ruan, J., Homer, N., Marth, G., Abecasis, G. & Durbin, R. The Sequence Alignment/Map format and SAMtools. *Bioinformatics* 25, 2078-2079 10.1093/bioinformatics/btp352 (2009).

46. McKenna, A., Hanna, M., Banks, E., Sivachenko, A., Cibulskis, K., Kernytsky, A., Garimella, K., Altshuler, D., Gabriel, S., Daly, M. & DePristo, M.A. The Genome Analysis Toolkit: a MapReduce framework for analyzing next-generation DNA sequencing data. *Genome Research* **20**, 1297-1303 10.1101/gr.107524.110 (2010).

47. Van der Auwera, G.A., Carneiro, M.O., Hartl, C., Poplin, R., Del Angel, G., Levy-Moonshine, A., Jordan, T., Shakir, K., Roazen, D., Thibault, J., Banks, E., Garimella, K.V., Altshuler, D., Gabriel, S. & DePristo, M.A. From FastQ data to high confidence variant calls: the Genome Analysis Toolkit best practices pipeline. *Current Protocols in Bioinformatics* **11**, 1110 10.1002/0471250953.bi1110s43 (2013).

48. Baasner, J.S., Howard, D. & Pucker, B. Influence of neighboring small sequence variants on functional impact prediction. *bioRxiv* posted 2019-06-13 10.1101/596718 (2019).

49. Schilbert, H.M., Pucker, B., Ries, D., Viehover, P., Micic, Z., Dreyer, F., Beckmann, K., Wittkop, B., Weisshaar, B. & Holtgrawe, D. Mapping-by-Sequencing Reveals Genomic Regions Associated with Seed Quality Parameters in Brassica napus. Genes 13, 10.3390/genes13071131 (2022).

50. Hackl, T. & Ankenbrand, M.J. gggenomes: A Grammar of Graphics for Comparative Genomics. R package version 0.9.5.9000 edn (2022).

51. Theine, J., Holtgrawe, D., Herzog, K., Schwander, F., Kicherer, A., Hausmann, L., Viehover, P., Topfer, R. & Weisshaar, B. Transcriptomic analysis of temporal shifts in berry development between two grapevine cultivars of the Pinot family reveals potential genes controlling ripening time. BMC Plant Biology 21, 327 10.1186/s12870-021-03110-6 (2021).

52. Arends, D., Prins, P., Jansen, R.C. & Broman, K.W. R/qtl: high-throughput multiple QTL mapping. *Bioinformatics* 26, 2990-2 10.1093/bioinformatics/btq565 (2010).

53. Koren, S., Walenz, B.P., Berlin, K., Miller, J.R., Bergman, N.H. & Phillippy, A.M. Canu: scalable and accurate long-read assembly via adaptive k-mer weighting and repeat separation. *Genome Research* **27**, 722-736 10.1101/gr.215087.116 (2017).

54. Bruna, T., Hoff, K.J., Lomsadze, A., Stanke, M. & Borodovsky, M. BRAKER2: automatic eukaryotic genome annotation with GeneMark-EP+ and AUGUSTUS supported by a protein database. NAR Genomics and Bioinformatics 3, lqaa108 10.1093/nargab/lqaa108 (2021).

55. Dobin, A., Davis, C.A., Schlesinger, F., Drenkow, J., Zaleski, C., Jha, S., Batut, P., Chaisson, M. & Gingeras, T.R. STAR: ultrafast universal RNA-seq aligner. Bioinformatics 29, 15–21 10.1093/bioinformatics/bts635 (2013).

56. Liao, Y., Smyth, G.K. & Shi, W. featureCounts: an efficient general purpose program for assigning sequence reads to genomic features. Bioinformatics 30, 923–930 10.1093/bioinformatics/btt656 (2014).

57. Love, M.I., Huber, W. & Anders, S. Moderated estimation of fold change and dispersion for RNA-seq data with DESeq2. Genome Biology 15, 550 10.1186/s13059-014-0550-8 (2014).

58. R Core Team. R: A language and environment for statistical computing. (R Foundation for Statistical Computing, Vienna, Austria, 2018).

59. Wickham, H. ggplot2: Elegant Graphics for Data Analysis (Springer, 2016).

60. Tang, H., Bowers, J.E., Wang, X., Ming, R., Alam, M. & Paterson, A.H. Synteny and collinearity in plant genomes. *Science* 320, 486-8 10.1126/science.1153917 (2008).

61. Altschul, S.F., Gish, W., Miller, W., Myers, E.W. & Lipman, D.J. Basic local alignment search tool. Journal of Molecular Biology 215, 403–410 10.1016/S0022-2836(05)80360-2 (1990).

62. Quevillon, E., Silventoinen, V., Pillai, S., Harte, N., Mulder, N., Apweiler, R. & Lopez, R. InterProScan: protein domains identifier. Nucleic Acids Research 33, W116–20 10.1093/nar/gki442 (2005).

63. Goel, M., Sun, H., Jiao, W.B. & Schneeberger, K. SyRI: finding genomic rearrangements and local sequence differences from whole-genome assemblies. Genome Biol 20, 277 10.1186/s13059-019-1911-0 (2019).

64. Mi, H., Muruganujan, A., Ebert, D., Huang, X. & Thomas, P.D. PANTHER version 14: more genomes, a new PANTHER GO-slim and improvements in enrichment analysis tools. Nucleic Acids Research 47, D419–D426 10.1093/nar/gky1038 (2019).

65. Katoh, K. & Standley, D.M. MAFFT multiple sequence alignment software version 7: improvements in performance and usability. Molecular Biology and Evolution 30, 772–780 10.1093/molbev/mst010 (2013).

