## Additional file S2 for "Genomic characterization of a nematode tolerance locus in sugar beet"

**S2A: Histogram of adjusted cyst counts (SN) data per family. SN = squared root of the number of cysts counted per plant (defined as in the methods section).**

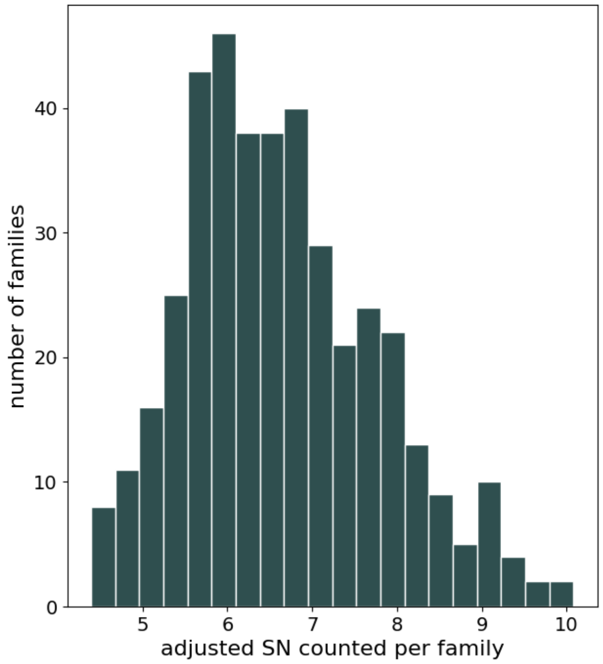

**S2B: Structural rearrangements between 2320BvONT and U2BvONT as identified with SyRI.**

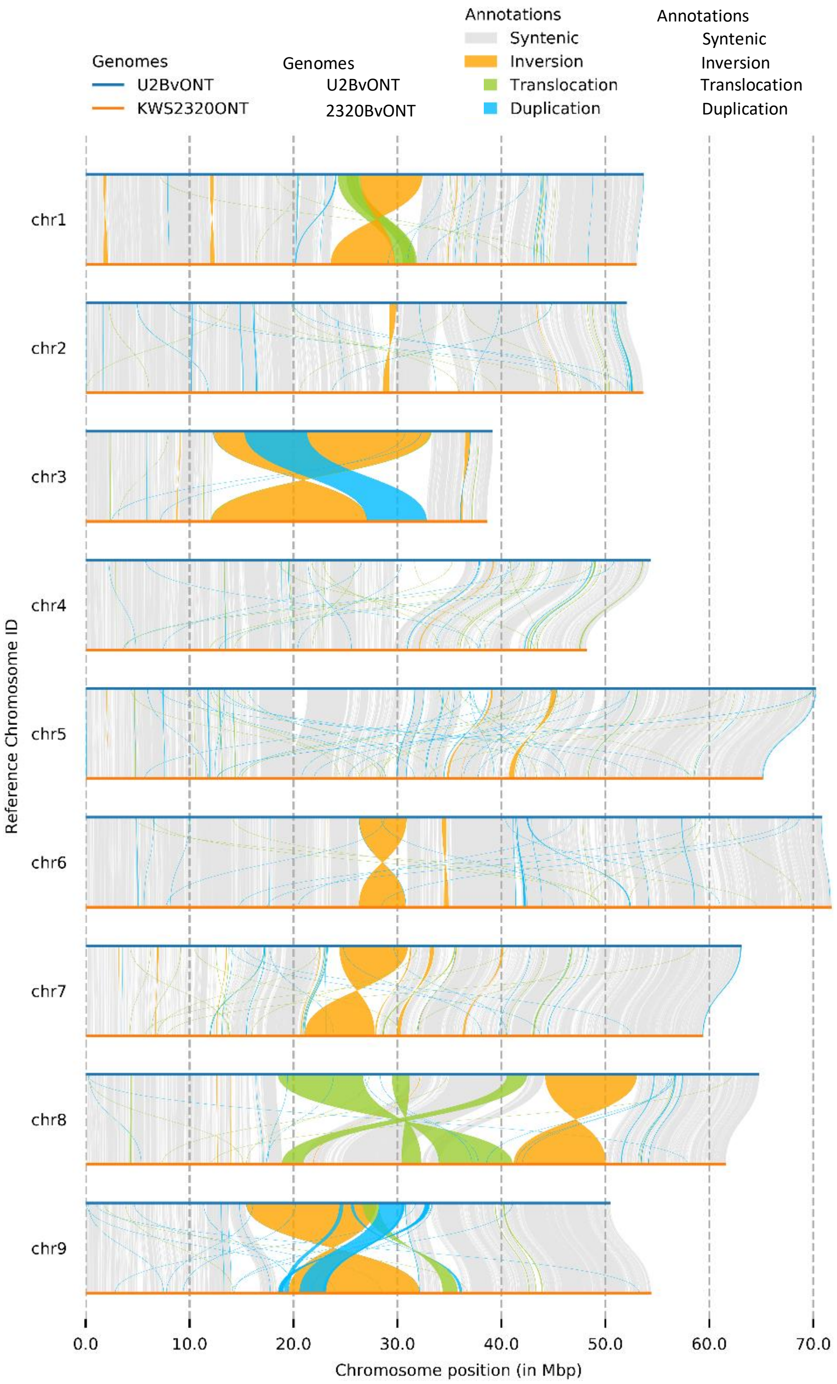

**S2C: Delta allele frequency plots for 10 SNP windows of all nine U2BvONT pseudochromosomes.**

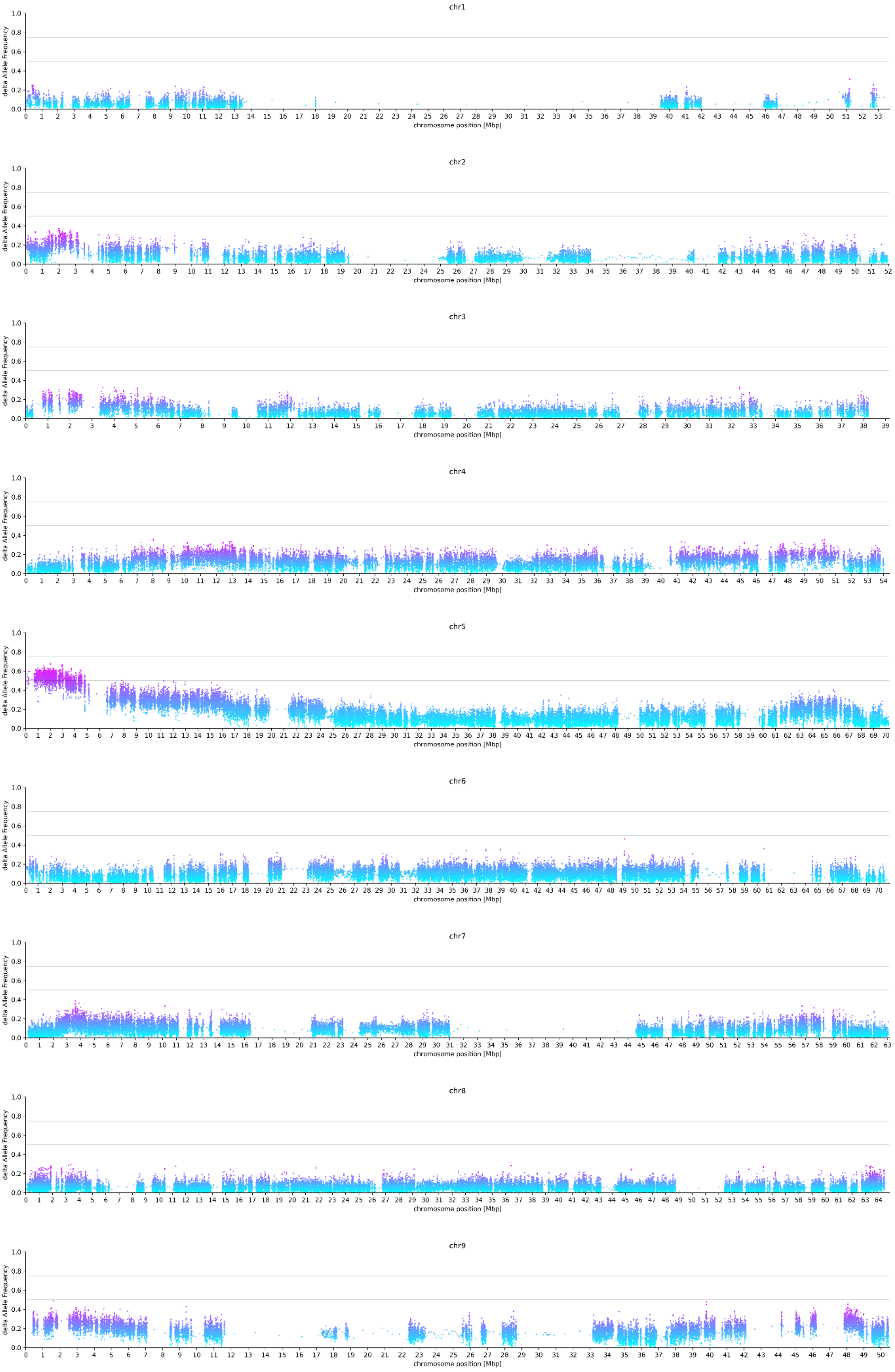

**S2D: Top: marker information/Primer sequences used for the determination of the potential NT locus. Bottom: graphical genotyping with flanking and co-segregating markers. Only data for informative genotypes/recombinants are shown.**

| Marker name | Type – Detection | Primer name | Sequence | Gene region | U2Bv-ONT chr05 position |
|---|---|---|---|---|---|
| MH00/01 | SNV-Sanger sequencing | MH00 | AGTCTCCGACTGCT CTATCTCC | Bv05_g11417_rnnp | 1,321,375 |
| MH00/01 | SNV-Sanger sequencing | MH01 | AAGCCTAAAGGAG GCTCTAACC | Bv05_g11417_rnnp | 1,322,283 |
| Bv5_g02 | SNV-Sanger sequencing | JG02 | AAGGAGCAAAATCT TCATTGAG | Bv05_g11456_dknx | 1,747,928 |
| Bv5_g02 | SNV-Sanger sequencing | JG01 | TCATAGCTTTCATA TCCCAAATTC | Bv05_g11456_dknx | 1,748,838 |
| Bv5_g05 | SNV-Sanger sequencing | JG07 | TTCTCTGATGATGA AAGCAATG | Bv05_g11459_hany | 1,813,501 |
| Bv5_g05 | SNV-Sanger sequencing | JG08 | TCTCCTTCTTCTGT CTCTTTCG | Bv05_g11459_hany | 1,814,483 |
| Bv5_g19 | SNV-Sanger sequencing | JG05 | TTGATTTCTTCAAAT GGGTTTC | Bv05_g11472_qkha | 1,910,354 |
| Bv5_g19 | SNV-Sanger sequencing | JG06 | AGGATCCTTCTTCC TACAAAGC | Bv05_g11472_qkha | 1,910,874 |
| BR1180 | SNV-KASPar | Bv05_BR1180-allele 1 | GACCATTCATAAGA AGCTCATCTTCA | Bv05_g11478_haee | 2,021,918 |
| BR1180 | SNV-KASPar | Bv05_BR1180-allele 2 | GACCATTCATAAGA AGCTCATCTTCG | Bv05_g11478_haee | 2,021,918 |
| BR1180 | SNV-KASPar | Bv05_BR1180-common primer | GAAGGGTCACCTCT CATTGCCATTA | Bv05_g11478_haee | 2,021,970 |
| TK15/16 | 341 bp InDel – agarose gel | TK16 | GCAAGGATGCAGA AAAATTACCC | Bv05_g11509_zukd | 2,343,550 |
| TK15/16 | 341 bp InDel – agarose gel | TK15 | AGAGGAGCTGCTTA CTTTGC | Bv05_g11509_zukd | 2,344,178 |
| SNP192 | SNV-HRM | SNP192-fw | CAGGCTTGAGCTGT TTGGCTATATA | - | 2,700,329 |
| SNP192 | SNV- HRM | SNP192-rev | CCTATAACGCATCT AAATGACAGGGT | - | 2,700,363 |

| | | | **Marker names ordered by physical position on U2Bv chromosome 5** | | | | | | |
|---|---|---|---|---|---|---|---|---|---|
| **Genotype** | **Description** | **Phenotype** | **MH00/01** | **Bv5_g02** | **Bv5_g05** | **Bv5_g19** | **BR1180** | **TK15/16** | **SNP192** |
| KWS2320 | sus. reference genotype | NS | A | A | A | A | A | A | A |
| U1Bv | sus. parental genotype (P1) | NS | A | A | A | A | A | A | A |
| U2Bv | tol. parental genotype (P2) | NT | B | B | B | B | B | B | B |
| F1 | F1 genotype used to generate F2 | | H | H | H | H | H | H | H |
| SUS3 | sus. standard line 3 | NS | A | A | A | A | A | A | A |
| BR12 | F2 genotype | NT | B | B | B | B | B | B | B |
| BR37 | F2 genotype | NT | B | B | B | B | B | B | B |
| BR38 | F2 genotype | NT | H | B | B | B | B | B | B |
| BR46 | F2 genotype | NS | H | H | A | A | A | A | A |
| BR47 | F2 genotype | NS | A | A | A | A | H | H | H |
| BR49 | F2 genotype | NS | A | A | A | A | H | H | H |
| BR50 | F2 genotype | NS | A | A | A | A | H | H | H |
| BR51 | F2 genotype | NS | B | H | H | H | H | H | H |

Abbreviations: sus. = susceptible; tol. = tolerant; NS = nematode susceptible; NT = nematode tolerant

**S2E: Visualization of key polymorphisms at conserved positions of Arabidopsis NLP7.**

S205

| | | |
|---|---|---|
| *Arabidopsis* NLP7 | L--GQPFVLNPNGNGL-NQYRMI**S**LTYMFSVDSESDVELGLPGR | |
| NLP7-T1 (Bv05_g11459_hany.t1) | NYHRQPFVKSLHLQFLDDEFGDL**K**LLKELT----HVTHLYISGD | lysine |
| NLP7-T2 (Bv05_g11461_tlqs.t1) | NYDFQPSVKFLDLNFLDDEFGDL**T**LLKELT----HVTHLYISGH | other polar aa and possible phosphorylation site |
| NLP7-T3 (Bv05_g11464_mdsr.t1) | NYDFQPSVKFLDLNFLDDEFGDL**T**LLKELT----HVTHLYISGH | other polar aa and possible phosphorylation site |
| NLP7-T4 (Bv05_g11465_rycp.t1) | S--GQPFVLGPHSNGL-NQYRTV**S**LMYMFSVDGESVTTLGLPGR | conserved |
| NLP7-S1 (Bv05_g11045_mdsr.t1) | NYDFQTSVKFLDLQFLDDEFGDL**T**LLKELT----HVTHLYISSN | other polar aa and possible phosphorylation site |
| NLP7-S2 (Bv05_g11050_nxxw.t1) | ------------------------------------------- | truncated |
| NLP7-S3 (Bv05_g11053_rycp.t1) | S--GQPFVLGPHSNGL-NQYRTV**S**LMYMFSVDGESVTTLGLPGR | conserved |

K867 D909 D911 E913

| | | |
|---|---|---|
| *Arabidopsis* NLP7 | SGSEMRTVTI**K**ASYKDDIIRFRISSGSGIMELKDEVAKRLKVDAGTFDIKYL**D**D**D**N**E**WVLIACDADL | |
| NLP7-T1 (Bv05_g11459_hany.t1) | SP-SLSKFVV**K**ATYKDDDYRIELPSTASFHELKKEVARTLELEMCKFKIRYK**D**E**D**N**E**WTRMSIDSHL | all conserved |
| NLP7-T2 (Bv05_g11461_tlqs.t1) | SP-SLSKIIV**K**ATYKDDTVRFHISSTASFHELEKEVAKNLKLKLSKFKIKYQ**D**E**D**K**E**LMLMTLDSHL | all conserved |
| NLP7-T3 (Bv05_g11464_mdsr.t1) | ------------------------------------------------------------------ | completely missing |
| NLP7-T4 (Bv05_g11465_rycp.t1) | ---DMKTVTI**K**ATFREDIIRFRFSLNSNIVELKEEVAKRFKLEVGTFEIKYL**D**D**D**H**E**WVLIACDSDL | all conserved |
| NLP7-S1 (Bv05_g11045_mdsr.t1) | SP-SLSKIIV**K**ATYKDDTVRFHIFSTTSFHELEKEVAKNLKLKLSKFKIKYE**D**E**E**K**E**LILMTLDSHL | all but D911 |
| NLP7-S2 (Bv05_g11050_nxxw.t1) | SP-NLSKLKV**K**ATYKKDDYSIELSSTPSFHELEKGVAQTIKLEMGKFKIKYK**D**E**D**N**E**WMRMTIDFHV | all conserved |
| NLP7-S3 (Bv05_g11053_rycp.t1) | ---DMKTVTI**K**ATFREDIIRFRFSLSSNIVELKEEVAKRFKLEVGTFEIKYL**D**D**D**H**E**WVLIACDSDL | all conserved |

**S2F: Principal component analysis for all samples of the RNA-Seq infection experiment.**

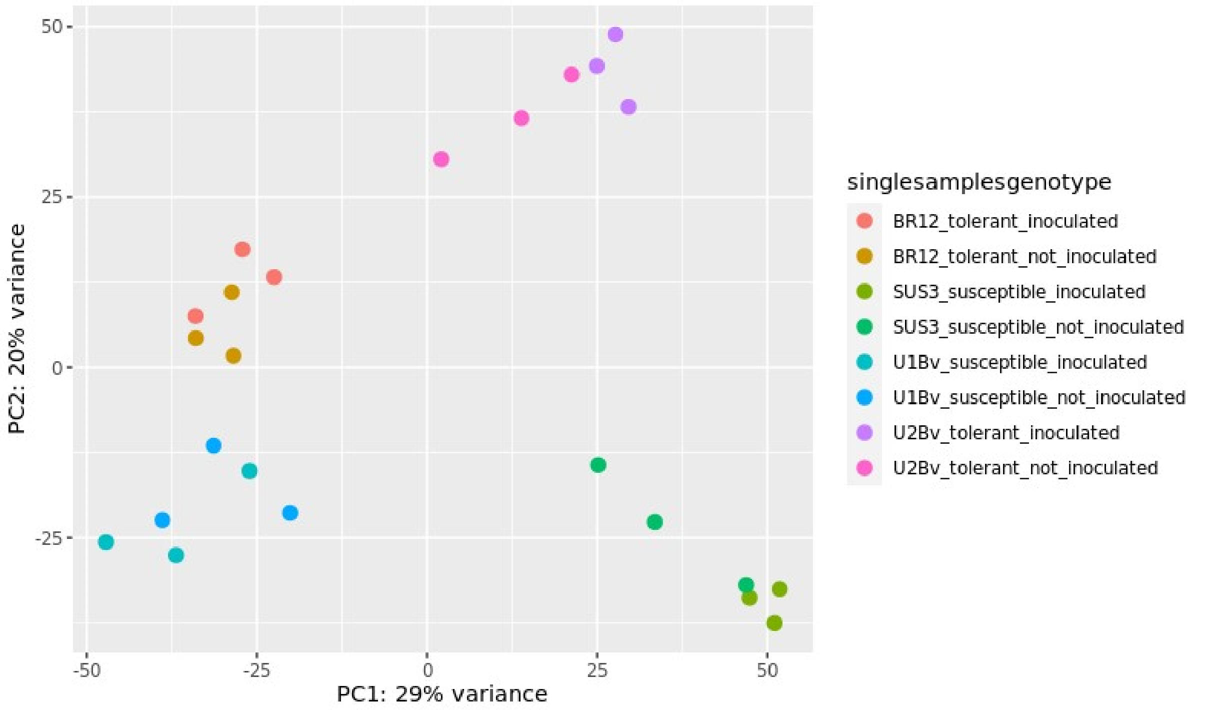

**S2G: Availability of datasets generated for this study.**

| data type | genotype | Source (SRA IDs/ENA IDs/DOI) |
|---|---|---|
| Mapping-by-sequencing Illumina reads | Strube U2Bv | PRJEB56338 - runs ERR10772555 - ERR10772680 |
| ONT reads | Strube U2Bv | PRJEB37059 |
| ONT reads | 2320Bv | PRJEB36905 |
| Illumina polishing reads | Strube U2Bv | PRJEB56338 - runs ERR10687994, ERR10687995 |
| Illumina polishing reads | 2320Bv | SRR952955, SRR952954, SRR952950, SRR952949, SRR952948, SRR952946, SRR869632, SRR869633, SRR869628, SRR869627, SRR869626, SRR868941, SRR868940, SRR868937, SRR868938, SRR868936, SRR868935, SRR868934, SRR868933, SRR869631 |
| RNA-Seq reads infection assay | Strube U2Bv | PRJEB58360 - runs ERR10691654, ERR10691655, ERR10691656, ERR10691657, ERR10691658, ERR10691659, ERR10691660, ERR10691661, ERR10691662, ERR10691663, ERR10691664, ERR10691665, ERR10691666, ERR10691667, ERR10691668. ERR10691669, ERR10691670, ERR10691671, ERR10691672, ERR10691673, ERR10691674, ERR10691675, ERR10691676, ERR10691677, ERR10691678, ERR10691679, ERR10691680, ERR10691681, ERR10691682, ERR10691683, ERR10691684, ERR10691685, ERR10691686, ERR10691687, ERR10691688, ERR10691689, ERR10691690, ERR10691691, ERR10691692, ERR10691693, ERR10691694, ERR10691695, ERR10691696, ERR10691697, |

| | | |
|---|---|---|
| | | ERR10691698, ERR10691699, ERR10691700, ERR10691701, ERR10691702, ERR10691703, ERR10691704, ERR10691705, ERR10691706, ERR10691707, ERR10691708, ERR10691709, ERR10691710, ERR10691711, ERR10691712, ERR10691713, ERR10691714, ERR10691715, ERR10691716, ERR10691717, ERR10691718, ERR10691719, ERR10691720, ERR10691721, ERR10691722, ERR10691723, ERR10691724, ERR10691725, ERR10691726, ERR10691727, ERR10691728, ERR10691729, ERR10691730, ERR10691731, ERR10691732, ERR10691733, ERR10691734, ERR10691735, ERR10691736, ERR10691737, ERR10691738, ERR10691739, ERR10691740, ERR10691741, ERR10691742, ERR10691743, ERR10691744, ERR10691745, ERR10691746, ERR10691747, ERR10691748, ERR10691749 |
| Assembly U2BvONT v1.0 | Strube U2Bv | PRJEB56338 - accession ERZ15181926 |
| Assembly 2320BvONT v1.0 | 2320Bv | PRJEB56338 - accession ERZ15181925 |
| Structural annotation, functional annotation and mRNA coverage/length files | Strube U2Bv | 10.4119/unibi/2966927 |
| Structural annotation, functional annotation and mRNA coverage/length files | 2320Bv | 10.4119/unibi/2966927 |

**S2H: Detailed methods for i) calculation of the delta allele frequencies and interval detection, ii) ONT sequencing, iii) ONT assembly, and iv) gene prediction and functional annotation.**

**i)** VCF files of individual samples were merged with a customized Python script to generate a multi-sample VCF file (VCF_combiner.py, https://github.com/bpucker/beetresmabs). This set of sequence variants in the F2 population was filtered based on variants detected in the F1 and the initial parents against the same reference sequence. The filtering was performed by the generation of a set of trusted variants ('gold standard'). A customized Python script (filter_parent_variants.py) was applied to keep only biallelic variants with a heterozygous state in the F1 and contrasting homozygous states in the initial parents. Variants were considered heterozygous if the alternative allele frequency was between 0.3 and 0.7. The classification '1/1' provided by GATK was used as flag to identify homozygous variants. Next, this set of sequence variants ('gold standard') served as basis to filter the variants detected in the F2 population with a customized Python script (filter_VCF_by_goldstandard.py). Only surviving variants in the F2 population were subjected to the dAF analysis that was performed for contigs longer than 1 Mbp with a customised Python script (dAF_selected_contigs.py). Two types of figures are generated: i) dAF values of individual variants are displayed based on their chromosomal location and ii) average dAF values of a certain number of adjacent variant positions are displayed. Default options for the second approach merge the values of 10 adjacent variant positions and shift the window of considered variant positions by 5 positions. Visualization of the results is based on the Python module matplotlib [1]. Based on the visualized dAF frequencies, the NT locus for BCN tolerance was identified. As a cutoff, we looked for a region with a dAF > 0.5 throughout five consecutive 10 SNP windows (50 bp). A dAF > 0.5 was only detected on chr5 (between positions 109,475 bp - 4,822,242 bp) indicating that only one candidate region is present. The NT locus was further restricted using genetic markers. BR1180 enabled the determination of a border in the south, where the SNP position in U2BvONT is at 2,021,943 bp. MH00/01 was used to determine the border in the north. The primer sequences and further information for both markers are available in Additional file S2D.

**ii)** High molecular weight genomic DNA of the nematode-tolerant parent of the mapping population Strube U2Bv and the sugar beet reference genotype 2320Bv was subjected to quality control via agarose gel electrophoresis, NanoDrop measurement, and Qubit measurement as previously described [2]. Short DNA fragments were depleted with the short read eliminator kit (Circulomics). ONT sequencing libraries were prepared following the LSK109 protocol. Sequencing was performed on seven R9.4.1 flow cells for Strube U2Bv and six flow cells (R9.4.1 and R10) for 2320Bv, respectively, on a GridION at the Sequencing Core Facility of the Center for Biotechnology (Bielefeld, Germany). Guppy v3 (https://nanoporetech.com/) was used for real time basecalling on the GridION. Sequencing output of the flow cells was increased by nuclease flushes upon demand to recover blocked pores as previously described [2]. FAST5 and FASTQ files were submitted to the European Nucleotide Archive (Additional file S2G).

For the U2BvONT and 2320BvONT genome sequence assembly, 43 Gbp and 50 Gbp of ONT sequencing read data with a read N50 length of 29 kbp and 28 kbp, respectively, were generated.

**iii)** Long reads of the nematode-tolerant parent (Strube U2Bv) and the sugar beet reference genotype 2320Bv were subjected to *de novo* genome assembly with Canu v1.8 [3]. The genome sequences of both genotypes were assembled with the following parameters: 'genomeSize=750m', 'corMhapFilterThreshold=0.0000000002', 'ovlMerThreshold=500', and 'corMhapOptions=--threshold 0.80 --num-hashes 512 --num-min-matches 3 --ordered-sketch-size 1000 --ordered-kmer-size 14 --min-olap-length 2000 --repeat-idf-scale 50'. Polishing of the initial assemblies with ONT long reads was performed with racon [4] and medaka (https://github.com/nanoporetech/medaka) as previously described [2]. Short read polishing was performed with three rounds of pilon [5] as previously described [6].

The final 2320BvONT v1.0 assembly was used to bridge two contigs of the initial Strube U2Bv assembly. Alignment of these two contigs to a single contig in the 2320BvONT v1.0 assembly revealed a gap of approx. 200 kbp between two contigs, which was closed by a Python script through long read walking (LRW.py, https://github.com/bpucker/LongReadWalker). The terminal 3 kbp of an existing contig were used as bait sequence to identify all matching ONT long reads via blastn v2.8.1 [7]. Filter criteria for suitable BLAST hits are at least 80% sequence similarity and a minimal alignment length of 1 kbp. The read with the best BLAST hit is used to extend the existing contig. An iterative search for matching reads was continued for 20 rounds with 3 kbp of the new terminal sequence. This gap closing approach was started from both contig ends and resulted in an overlapping sequence. The genes dknx and orwr are the last genes on the two former contigs which were connected by ‘long read walking’ (Figure 2). Finally, the gap filling sequence was improved through the assembly polishing process (Additional file S2K).

After the polishing process, contigs below a length of 100 kbp were discarded. Further, as previously described [2], contigs with matches to a black list were discarded in the ‘decontamination’ process. Scaffolding was performed with ALLMAPS v1.2.4 [8] using genetic markers for anchoring. First, the action ‘merge’ was used to merge maps in bed format, followed by the actual scaffolding performed with the action ‘path’. Both actions were performed with default parameters.

**iv)** Repeat masking of both genome assembly sequences was performed to get an overview of the repeat content in both genomes and to enhance gene prediction accuracy during the annotation process [9]. First, a *de novo* species-specific repeat library was constructed using RepeatModeler v2.0 [10]. RepeatModeler was used with the integrated LTR discovery pipeline. The resulting library, together with the Beta vulgaris repeat database retrieved from RepBase [11], was used as input for softmasking with RepeatMasker v4.1.1 [12].

As input for the BRAKER2 pipeline used for gene prediction [9], protein evidence and RNA hints were generated.

Protein evidence was retrieved from OrthoDB protein sequences [13] and hint files were generated with ProtHint [14].

In the next step, full length sugar beet mRNAs from previous annotations (RefBeet1.0 [15] and BeetSet2 [16] (including minor manual curated improvements)) were prepared to be integrated into the annotation pipeline. The mRNAs were aligned to the genome assembly sequences via BLAT [17] with the options '-fine' and '-q=rna'. Then, the alignments were filtered using the perl script 'filterPSL.pl' to output all best matches (--best, --minCover=80, --minId=92). After sorting by sequence names and by start coordinates, 'blat2hints.pl' was applied to the alignment files to generate the hint files in GFF format. The hint files derived from RefBeet1.0 and BeetSet2 were compared by position of the alignments and in case of an overlap of two alignments, the hit of the respective BeetSet mRNA was used for the final merged hints file.

In addition, RNA-Seq data from various conditions and tissues was used to improve the gene prediction (Additional file S2J). First, the reads were mapped to the genome assembly sequences with STARlong v2.7.6a [18]. For Strube U2Bv, the RNA-Seq reads were mapped to a combined reference including the genome assembly sequence of *H. schachtii* (NCBI accession PRJNA722882). Mappings onto the nematode genome sequence were discarded. Then, primary alignments on the U2BvONT sequence or the 2320BvONT sequence, respectively, were extracted and the AUGUSTUS script 'bam2hints' was applied to generate hints in GFF format.

As recommended in the BRAKER2 documentation, AUGUSTUS prediction with BRAKER2 was run twice per genome assembly - separately with protein (OrthoDB) and RNA evidence (full length mRNAs from previous annotations and RNA-Seq data). The results of both runs were combined with TSEBRA [19].

As a next part of the annotation pipeline, gene model refinement was performed with PASA [20]. To generate a suitable input for PASA, Trinity v2.12.0 [21,22] was used with default parameters for genome-guided *de novo* transcriptome assemblies. The script 'Launch_PASA_pipeline.pl' was then used with default parameters providing the respective transcriptome assembly as input to align transcripts to the genome assembly sequences. In the next step, annotation comparison was performed to generate an updated gene set. Two rounds of PASA were performed for each genome assembly/annotation. tRNAs were predicted using tRNA Scan-SE [23] as submodule of the 'RNA Gatherer' pipeline and then merged with the final PASA gene prediction.

The gene models were further filtered. Identical gene models (same gene name, same CDS and same protein sequence, but alternative PASA models) and genes with an amino acid length smaller than 50 were discarded.

The gene models within the respective NT loci were manually refined using IGV v2.8.0 [24] and WebApollo Annotator [25]. Previously published full length cDNAs, RNASeq coverage and splice junction information derived from the BAM files as well as RefBeet gene models were used for visual input and comparison. Liftoff v1.6.1 [26] was used to transfer the RefBeet/BeetSet2 annotation to the respective new genome coordinates. The U2BvONT annotation was also transferred to the new 2320BvONT reference sequence coordinates and *vice versa*. The BAM files were prefiltered to only include mappings with a phred score of 50 or higher. In most cases, UTR sequences were added to the existing gene models according to available RNA-Seq coverage. Additionally, transcript variants and gene models from the other respective genome assembly annotation and from the RefBeet annotation were added if the resulting transcripts had a minimal length of 50 amino acids or 150 base pairs and if coverage and splice junction support by RNA-Seq data was present.

New gene names were assigned to all final gene models. If a gene-based RBH with the RefBeet annotations was identified, the same four letter code was integrated into the naming process. Otherwise, a random new combination of four letters was assigned to the respective gene model. Each gene was named according to the following convention: starting with the species abbreviation 'Bv', followed by the chromosome number (e.g. Bv01), followed by the gene number in the assembly based on ascending coordinates (e.g. Bv01_g1) and finally followed by the four letter code (e.g. Bv01_g1_zjcg).

A functional annotation was constructed for the final gene prediction. Therefore, the results of a InterProScan (v5.52) [27] run, a SwissProt BLASTX [7] and a RBH BLAST against the published RefBeet annotations, were combined in a single file (available at DOI 10.4119/unibi/2966927).

For each transcript, the percentage of coverage over the transcript length and the average coverage were calculated (available at DOI 10.4119/unibi/2966927). This enables the identification of high and low confidence gene models.

**References**


1. Hunter, J.D. Matplotlib: A 2D Graphics Environment. *Comput. Sci. Eng.* **2007**, *9*, 90–95, doi:10.1109/MCSE.2007.55.
2. Siadjeu, C.; Pucker, B.; Viehöver, P.; Albach, D.C.; Weisshaar, B. High Contiguity de Novo Genome Sequence Assembly of Trifoliate Yam (Dioscorea Dumetorum) Using Long Read Sequencing. *Genes* **2020**, *11*, 274, doi:10.3390/genes11030274.
3. Koren, S.; Walenz, B.P.; Berlin, K.; Miller, J.R.; Bergman, N.H.; Phillippy, A.M. Canu: Scalable and Accurate Long-Read Assembly via Adaptive *k* -Mer Weighting and Repeat Separation. *Genome Res.* **2017**, *27*, 722–736, doi:10.1101/gr.215087.116.
4. Vaser, R.; Sović, I.; Nagarajan, N.; Šikić, M. Fast and Accurate de Novo Genome Assembly from Long Uncorrected Reads. *Genome Res.* **2017**, *27*, 737–746, doi:10.1101/gr.214270.116.
5. Walker, B.J.; Abeel, T.; Shea, T.; Priest, M.; Abouelliel, A.; Sakthikumar, S.; Cuomo, C.A.; Zeng, Q.; Wortman, J.; Young, S.K.; et al. Pilon: An Integrated Tool for Comprehensive Microbial Variant Detection and Genome Assembly Improvement. *PLoS ONE* **2014**, *9*, e112963, doi:10.1371/journal.pone.0112963.
6. Pucker, B.; Holtgräwe, D.; Stadermann, K.B.; Frey, K.; Huettel, B.; Reinhardt, R.; Weisshaar, B. A Chromosome-Level Sequence Assembly Reveals the Structure of the Arabidopsis Thaliana Nd-1 Genome and Its Gene Set. *PLOS ONE* **2019**, *14*, e0216233, doi:10.1371/journal.pone.0216233.
7. Altschul, S.F.; Gish, W.; Miller, W.; Myers, E.W.; Lipman, D.J. Basic Local Alignment Search Tool. *J. Mol. Biol.* **1990**, *215*, 403–410, doi:10.1016/S0022-2836(05)80360-2.
8. Tang, H.; Zhang, X.; Miao, C.; Zhang, J.; Ming, R.; Schnable, J.C.; Schnable, P.S.; Lyons, E.; Lu, J. ALLMAPS: Robust Scaffold Ordering Based on Multiple Maps. *Genome Biol.* **2015**, *16*, 3, doi:10.1186/s13059-014-0573-1.
9. Brůna, T.; Hoff, K.J.; Lomsadze, A.; Stanke, M.; Borodovsky, M. BRAKER2: Automatic Eukaryotic Genome Annotation with GeneMark-EP+ and AUGUSTUS Supported by a Protein Database. *NAR Genomics Bioinforma.* **2021**, *3*, lqaa108, doi:10.1093/nargab/lqaa108.
10. Flynn, J.M.; Hubley, R.; Goubert, C.; Rosen, J.; Clark, A.G.; Feschotte, C.; Smit, A.F. RepeatModeler2 for Automated Genomic Discovery of Transposable Element Families. *Proc. Natl. Acad. Sci.* **2020**, *117*, 9451–9457, doi:10.1073/pnas.1921046117.
11. Jurka, J.; Kapitonov, V.V.; Pavlicek, A.; Klonowski, P.; Kohany, O.; Walichiewicz, J. Repbase Update, a Database of Eukaryotic Repetitive Elements. *Cytogenet. Genome Res.* **2005**, *110*, 462–467, doi:10.1159/000084979.
12. Smit, A.; Hubley, R.; Green, P. RepeatMasker Open-4.0. 2015.

13. Kriventseva, E.V.; Kuznetsov, D.; Tegenfeldt, F.; Manni, M.; Dias, R.; Simão, F.A.; Zdobnov, E.M. OrthoDB V10: Sampling the Diversity of Animal, Plant, Fungal, Protist, Bacterial and Viral Genomes for Evolutionary and Functional Annotations of Orthologs. *Nucleic Acids Res.* **2019**, *47*, D807–D811, doi:10.1093/nar/gky1053.
14. Brůna, T.; Lomsadze, A.; Borodovsky, M. GeneMark-EP+: Eukaryotic Gene Prediction with Self-Training in the Space of Genes and Proteins. *NAR Genomics Bioinforma.* **2020**, *2*, lqaa026, doi:10.1093/nargab/lqaa026.
15. Dohm, J.C.; Minoche, A.E.; Holtgräwe, D.; Capella-Gutiérrez, S.; Zakrzewski, F.; Tafer, H.; Rupp, O.; Sörensen, T.R.; Stracke, R.; Reinhardt, R.; et al. The Genome of the Recently Domesticated Crop Plant Sugar Beet (Beta Vulgaris). *Nature* **2014**, *505*, 546–549, doi:10.1038/nature12817.
16. Minoche, A.E.; Dohm, J.C.; Schneider, J.; Holtgräwe, D.; Viehöver, P.; Montfort, M.; Rosleff Sörensen, T.; Weisshaar, B.; Himmelbauer, H. Exploiting Single-Molecule Transcript Sequencing for Eukaryotic Gene Prediction. *Genome Biol.* **2015**, *16*, 184, doi:10.1186/s13059-015-0729-7.
17. Kent, W.J. BLAT---The BLAST-Like Alignment Tool. *Genome Res.* **2002**, *12*, 656–664, doi:10.1101/gr.229202.
18. Dobin, A.; Davis, C.A.; Schlesinger, F.; Drenkow, J.; Zaleski, C.; Jha, S.; Batut, P.; Chaisson, M.; Gingeras, T.R. STAR: Ultrafast Universal RNA-Seq Aligner. *Bioinformatics* **2013**, *29*, 15–21.
19. Gabriel, L.; Hoff, K.J.; Brůna, T.; Borodovsky, M.; Stanke, M. TSEBRA: Transcript Selector for BRAKER. *BMC Bioinformatics* **2021**, *22*, 566, doi:10.1186/s12859-021-04482-0.
20. Haas, B.J. Improving the Arabidopsis Genome Annotation Using Maximal Transcript Alignment Assemblies. *Nucleic Acids Res.* **2003**, *31*, 5654–5666, doi:10.1093/nar/gkg770.
21. Grabherr, M.G.; Haas, B.J.; Yassour, M.; Levin, J.Z.; Thompson, D.A.; Amit, I.; Adiconis, X.; Fan, L.; Raychowdhury, R.; Zeng, Q.; et al. Full-Length Transcriptome Assembly from RNA-Seq Data without a Reference Genome. *Nat. Biotechnol.* **2011**, *29*, 644–652, doi:10.1038/nbt.1883.
22. Haas, B.J.; Papanicolaou, A.; Yassour, M.; Grabherr, M.; Blood, P.D.; Bowden, J.; Couger, M.B.; Eccles, D.; Li, B.; Lieber, M.; et al. De Novo Transcript Sequence Reconstruction from RNA-Seq Using the Trinity Platform for Reference Generation and Analysis. *Nat. Protoc.* **2013**, *8*, 1494–1512, doi:10.1038/nprot.2013.084.
23. Chan, P.P.; Lowe, T.M. TRNAscan-SE: Searching for TRNA Genes in Genomic Sequences. In *Gene Prediction*; Kollmar, M., Ed.; Methods in Molecular Biology; Springer New York: New York, NY, 2019; Vol. 1962, pp. 1–14 ISBN 978-1-4939-9172-3.
24. Robinson, J.T.; Thorvaldsdóttir, H.; Winckler, W.; Guttman, M.; Lander, E.S.; Getz, G.; Mesirov, J.P. Integrative Genomics Viewer. *Nat. Biotechnol.* **2011**, *29*, 24–26, doi:10.1038/nbt.1754.
25. Lee, E.; Helt, G.A.; Reese, J.T.; Munoz-Torres, M.C.; Childers, C.P.; Buels, R.M.; Stein, L.; Holmes, I.H.; Elsik, C.G.; Lewis, S.E. Web Apollo: A Web-Based Genomic Annotation Editing Platform. *Genome Biol.* **2013**, *14*, R93, doi:10.1186/gb-2013-14-8-r93.
26. Shumate, A.; Salzberg, S.L. Liftoff: Accurate Mapping of Gene Annotations. *Bioinformatics* **2021**, *37*, 1639–1643, doi:10.1093/bioinformatics/btaa1016.
27. Quevillon, E.; Silventoinen, V.; Pillai, S.; Harte, N.; Mulder, N.; Apweiler, R.; Lopez, R. InterProScan: Protein Domains Identifier. *Nucleic Acids Res.* **2005**, *33*, W116–W120, doi:10.1093/nar/gki442.

**S2I: Overview of the RNA-Seq samples in the infection experiment.**

| Alias | Description | Accession |
|---|---|---|
| BvA | BR12 tolerant line inoculated replicate 1 | ERS14359733 |
| BvB | BR12 tolerant line inoculated replicate 2 | ERS14359734 |
| BvC | BR12 tolerant line inoculated replicate 3 | ERS14359735 |
| BvD | U1Bv susceptible line inoculated replicate 1 | ERS14359736 |
| BvE | U1Bv susceptible line inoculated replicate 2 | ERS14359737 |
| BvF | U1Bv susceptible line inoculated replicate 3 | ERS14359738 |
| BvG | U2Bv tolerant line inoculated replicate 1 | ERS14359739 |
| BvH | U2Bv tolerant line inoculated replicate 2 | ERS14359740 |
| BvI | U2Bv tolerant line inoculated replicate 3 | ERS14359741 |
| BvJ | SUS3 susceptible line inoculated replicate 1 | ERS14359742 |
| BvK | SUS3 susceptible line inoculated replicate 2 | ERS14359743 |
| BvL | SUS3 susceptible line inoculated replicate 3 | ERS14359744 |
| BvM | BR12 tolerant line not inoculated replicate 1 | ERS14359745 |
| BvN | BR12 tolerant line not inoculated replicate 2 | ERS14359746 |
| BvO | BR12 tolerant line not inoculated replicate 3 | ERS14359747 |
| BvP | U1Bv susceptible line not inoculated replicate 1 | ERS14359748 |
| BvQ | U1Bv susceptible line not inoculated replicate 2 | ERS14359749 |
| BvR | U1Bv susceptible line not inoculated replicate 3 | ERS14359750 |
| BvS | U2Bv tolerant line not inoculated replicate 1 | ERS14359751 |
| BvT | U2Bv tolerant line not inoculated replicate 2 | ERS14359752 |
| BvU | U2Bv tolerant line not inoculated replicate 3 | ERS14359753 |
| BvV | SUS3 susceptible line not inoculated replicate 1 | ERS14359754 |
| BvW | SUS3 susceptible line not inoculated replicate 2 | ERS14359755 |
| BvX | SUS3 susceptible line not inoculated replicate 3 | ERS14359756 |

**S2J: List of all RNA-Seq datasets incorporated as hints for the gene prediction.**

| Genotype | SRA IDs/ENA IDs |
|---|---|
| Strube U2Bv | ERS14360034, ERS14360035, ERS14360036, ERS14359739, ERS14359740, ERS14359741, ERS14359751, ERS14359752, ERS14359753 |
| 2320Bv | PRJEB58621 and PRJEB62793 - runs ERR11534264- ERR11534345<br><br>ERR048960, ERR2040223, ERR3452443, ERR3452444, ERR3452445, ERR3452446, ERR3452447, ERR3452448, ERR3452449, ERR3452450, ERR3452451, ERR3452452, ERR3452453, ERR3452454, ERR3452455, ERR3452456, ERR3452457, ERR3452458, ERR3452459, ERR3452460, ERR3452461, ERR3452462, ERR3452463, ERR3452464, ERR3452465, ERR3452466, ERR3452467, ERR3452468, ERR3452469, ERR3452470, ERR3452471, ERR3452472, ERR747962, SRR10037935, SRR10039075, SRR10039076, SRR10039077, SRR10039078, SRR10039079, SRR10039080, SRR10039081, SRR10039082, SRR10039083, SRR10039084, SRR10039085, SRR10039086, SRR10039087, SRR10039088, SRR10039089, SRR10039090, SRR10039091, SRR10039092, SRR10039093, SRR10039094, SRR10039095, SRR10039096, SRR10039097, SRR10039098, SRR10189432, SRR10189433, SRR10189434, SRR10189435, SRR10189436, SRR10189437, SRR1022496, SRR10990171, SRR10990172, SRR10990173, SRR10990174, SRR10990175, SRR10990176, SRR10990177, SRR10990178, SRR10990179, SRR10990180, SRR10990181, SRR10990182, SRR10990183, SRR10990184, SRR10990185, SRR10990186, SRR10990187, SRR10990188, SRR10990189, SRR10990190, SRR10990191, SRR10990192, SRR10990193, SRR10990194, SRR10990195, SRR10990196, SRR10990197, SRR10990198, SRR10990199, SRR10990200, SRR10990201, SRR10990202, SRR10990203, SRR10990204, SRR10990205, SRR10990206, SRR11243750, SRR11243751, SRR11243752, SRR11243753, SRR11243754, SRR11243755, SRR11243756, SRR11243757, SRR11243758, SRR11243759, SRR11243760, SRR11243761, SRR11243762, SRR11243763, SRR11243764, SRR11243765, SRR11243766, SRR11243767, SRR11243768, SRR11243769, SRR11243770, SRR11243771, SRR11243772, SRR11243773, SRR11243774, SRR11243775, SRR11243776, SRR11243777, SRR11243778, SRR11243779, SRR11243780, SRR11243781, SRR11243782, SRR11243783, SRR11243784, SRR11243785, SRR11243786, SRR11243787, SRR11243788, SRR11243789, SRR11243790, SRR11243791, SRR11243792, SRR11243793, SRR11243794, SRR11243795, SRR11243796, SRR11243797, SRR11243798, SRR11243799, SRR11243800, SRR11243801, SRR11243802, SRR11243803, SRR11243804, |

| | |
|---|---|
| | SRR11243805, SRR11243806, SRR11243807, SRR11243808, SRR11243809, SRR11243810, SRR11243811, SRR11243812, SRR11243813, SRR11243814, SRR11243815, SRR11243816, SRR11243817, SRR11243818, SRR11243819, SRR11243820, SRR11243821, SRR11243822, SRR11243823, SRR11243824, SRR11243825, SRR11243826, SRR11243827, SRR11243828, SRR11243829, SRR11243830, SRR11243831, SRR11243832, SRR11243833, SRR11243834, SRR11243835, SRR11243836, SRR11243837, SRR11243838, SRR11243839, SRR11243840, SRR11243841, SRR11243842, SRR11243843, SRR11243844, SRR11243845, SRR11821851, SRR11836729, SRR11836730, SRR11836731, SRR11836732, SRR11836733, SRR11836734, SRR11836735, SRR11836736, SRR11836737, SRR11836738, SRR11836739, SRR11836740, SRR11836741, SRR11836742, SRR11836743, SRR11836744, SRR11836745, SRR11836746, SRR11836747, SRR11836748, SRR11836749, SRR11836750, SRR11836751, SRR11836752, SRR11836753, SRR11836754, SRR11836755, SRR11836756, SRR11836757, SRR11836758, SRR12121649, SRR1508751, SRR1508753, SRR1508755, SRR1508756, SRR1508758, SRR1508782, SRR1542626, SRR1542627, SRR1699442, SRR3823656, SRR3823657, SRR3823689, SRR3823690, SRR3823691, SRR3823692, SRR4293384, SRR4293385, SRR4293386, SRR4293699, SRR4293700, SRR4294169, SRR4294172, SRR4294181, SRR4294193, SRR4294692, SRR5127351, SRR5127352, SRR5127353, SRR5127354, SRR5127355, SRR5127356, SRR5127357, SRR5127358, SRR5127359, SRR5127360, SRR5127361, SRR5127362, SRR5127363, SRR5127364, SRR5127365, SRR5127366, SRR5127367, SRR5127368, SRR5127369, SRR5127370, SRR5127371, SRR5127372, SRR5127373, SRR5127374, SRR5127375, SRR5127376, SRR5127377, SRR5127378, SRR5127379, SRR5127380, SRR5127381, SRR5127382, SRR5127383, SRR5127384, SRR5127385, SRR5127386, SRR6339627, SRR6339628, SRR6339629, SRR6339630, SRR6339631, SRR6339632, SRR6339633, SRR6339634, SRR6339635, SRR7152809, SRR7152810, SRR7152811, SRR7152812, SRR7152813, SRR7152814, SRR7152815, SRR7152816, SRR7152817, SRR7152818, SRR7152819, SRR7152820, SRR7152821, SRR7152822, SRR7152823, SRR7152824, SRR7152825, SRR7152826, SRR7152827, SRR7152828, SRR7152829, SRR7152830, SRR7152831, SRR7152832, SRR7152833, SRR7152834, SRR7152835, SRR7152836, SRR7152837, SRR7152838, SRR7152839, SRR7152840, SRR7152841, SRR7152842, SRR7152843, SRR7152844, SRR7152845, SRR7152846, SRR7152847, SRR7152848, SRR7152849, SRR7152850, SRR7152851, SRR7152852, SRR7152853, SRR7152854, SRR7152855, SRR7152856, SRR7152857, SRR7152858, SRR7152859, SRR7152860, SRR7152861, SRR7152862, SRR7152863, SRR7152864, SRR7152865, SRR7152866, SRR7152867, SRR7152868, SRR7226042, SRR7226043, SRR7226044, SRR7226045, SRR7226046, SRR7226047, SRR7226048, SRR7226049, SRR7226050, SRR7226051, SRR7226052, SRR7226053, SRR8088710, SRR8088711, SRR8088712, SRR8088713, SRR8088714, SRR8088715, SRR8510321, SRR8510322, SRR8510323, SRR8510324, SRR8510325, |

| | |
|---|---|
| | SRR8510326, SRR8510327, SRR8510328, SRR8510329, SRR8510330, SRR868803, SRR868804, SRR868805, SRR868886, SRR868887, SRR869750, SRR869751, SRR869752, SRR869753, SRR869754, SRR869755, SRR869756, SRR869757, SRR869758, SRR869759, SRR869760, SRR869761, SRR869762, SRR869763, SRR869764, SRR869765, SRR869766, SRR869767, SRR869768, SRR869769, SRR869770, SRR869771, SRR869772, SRR869773, SRR869774, SRR869775, SRR869776, SRR9925582, SRR9925583, SRR9925584, SRR9925585, SRR9925586, SRR9925587, SRR9925588, SRR9925589, SRR9925590, SRR9925591, SRR9925592, SRR9925593, SRR9925594, SRR9925595, SRR9925596, SRR9925597 |
| EL10 | SRR10039075, SRR10039076, SRR10039077, SRR10039078, SRR10039079, SRR10039080, SRR10039081, SRR10039082, SRR10039083, SRR10039084, SRR10039085, SRR10039086, SRR10039087, SRR10039088, SRR10039089, SRR10039090, SRR10039091, SRR10039092, SRR10039093, SRR10039094, SRR10039095, SRR10039096, SRR10039097, SRR10039098 |

**S2K: Dot plot heatmap of the closed gap region in the initial Strube U2Bv assembly**.

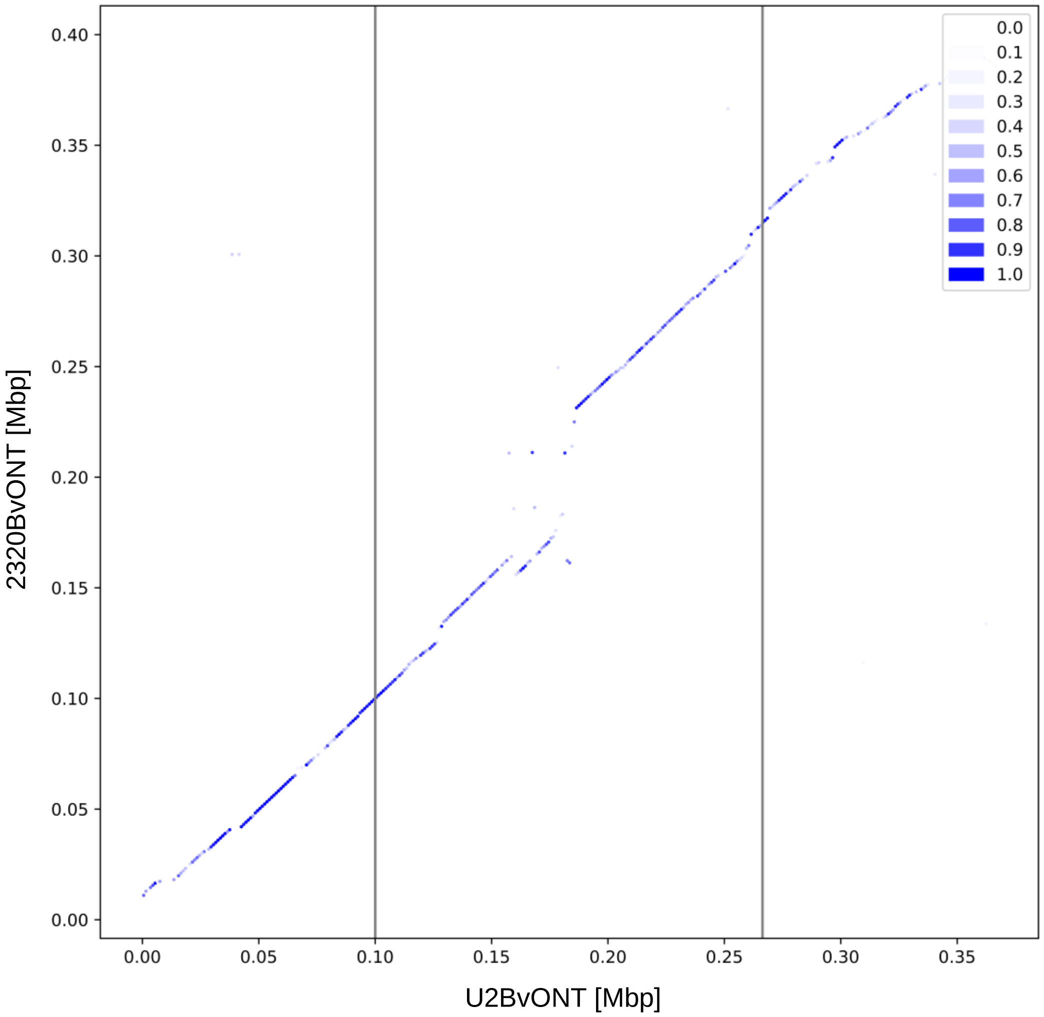


The gap between two contigs of the initial assembly of the genotype U2Bv was closed via long read walking. The vertical lines represent the respective borders (start/end) of the two former contigs. Starting from these border positions, the sequence was extended and the contigs were connected via long read walking. The part between the two vertical lines represents this new sequence. Dark blue dots show high sequence identity. The genes dknx and orwr are the last genes on the two former contigs which were connected by ‘long read walking’.
